# Spatial Logic Reconciles Gene-signature Methods in Triple Negative Breast Cancer

**DOI:** 10.64898/2026.08.13.744259

**Authors:** William Bastian, Jane L. Meisel, Ji-Hoon Lee, Nada Shaker, Lyra Griffiths, Meili Aiello, Zachary Buchwald, Yi Liu, E. Aubrey Thompson, Zaibo Li, Eugene F. Douglass, Xiaoxian Li

## Abstract

Triple-Negative Breast Cancer (TNBC) presents a significant clinical challenge due to its heterogeneity and lack of targeted treatment options, with chemotherapy and immunotherapy combinations currently serving as the main therapeutic strategy. Efforts to address TNBC heterogeneity have largely focused on classifying intrinsic cancer subtypes based on differential tumor mRNA expression, a strategy that has proven effective in hormone receptor-positive breast cancers but has yet to yield a clinically useful predictor of survival or treatment response in TNBC. We hypothesize that both the intrinsic characteristics of TNBC and the surrounding immune microenvironment influence treatment outcomes and that immune cell infiltration affects TNBC subtype classification and response variability.

To explore this hypothesis, we compared the predictive and prognostic capabilities of cancer subtype-based (TNBC-type) gene signatures and immune cell deconvolution methods (CIBERSORT) within the same TNBC datasets. We found that immune cell abundance outperformed TNBC subtype-signatures and multicellular immune cell aggregates showed the highest performance of all. More specifically, aggregate immune cells associated with tertiary lymphoid structures and tumor associated macrophages/monocytes demonstrated statistically significant predictive value. These findings were confirmed in an independent cohort of 67 TNBC patients treated with neoadjuvant chemotherapy. Further, single-cell RNA sequencing analysis revealed that the predictive power of cancer-subtype could be partially explained by immune- and stromal features. Examination of single-cell resolution spatial transcriptomic data confirmed presence of TLS-like, TAM- and cancer-stromal niches within TNBC biopsy samples that were associated with treatment response.

Overall, our results highlight that immune cell aggregates, which capture the spatial organization of the TME, outperform cell-type specific gene signatures in predicting TNBC outcomes. Our novel approach provides a robust framework for interpreting spatial relationships in bulk RNA-seq data, offering a pathway for reconciling past data with current advancements in spatial profiling technologies. This work paves the way for future studies to leverage the multi-cellular complexity of TNBC, enhancing diagnostic precision and facilitating the development of therapies that strategically modulate the tumor microenvironment for improved anti-cancer responses.

## INTRODUCTION

Triple-Negative Breast Cancer (TNBC) is a uniquely challenging and heterogeneous group of cancers defined by the absence of estrogen receptor (ER), progesterone receptor (PR), and HER2 expression.^1^ This lack of defined oncogenic drivers makes TNBC difficult to treat with targeted therapies, positioning chemotherapy and immunotherapy (chemo-IO) combinations as the primary treatment strategies, particularly in neoadjuvant and metastatic settings. Recent clinical trials have shown that chemo-IO can yield impressive pathological complete response (pCR) rates of up to 60%; however, patients with residual disease post-treatment remain at significantly elevated risk, with a six-fold increase in recurrence likelihood and twelve-fold increase in mortality compared to those achieving pCR.^2–4^ These contrasting outcomes highlight an urgent need for improved methods to define TNBC subtypes that are most likely to respond to chemo-IO regimens.

While chemo-IO like Keynote-522 (pembrolizumab + chemotherapy) represent a major advance, they are used without predictive biomarkers, leading to widespread overtreatment and avoidable toxicity^5–7^. At Emory, real-world data show similar pCR rates for KN522 (48.4%) and chemotherapy alone (47.4%), suggesting that many patients may not require immunotherapy^8^. Our outcomes are consistent with the high variability observed across chemoIO studies, where the addition of checkpoint therapy improves pCR rates by only 2.8–17% depending on the patient cohort.^9^ This has renewed interest in classifiers that can predict pCR for chemotherapy alone to guide de-escalation strategies.^5–7^ Critically, correlates—such as TNBC subtype and infiltrating immune cells— currently lack the resolution to guide therapy decisions.

### TNBC-subtypes

To address TNBC’s inherent heterogeneity, research efforts have increasingly focused on defining intrinsic cancer subtypes through RNA-seq-based multi-gene classifiers.^10–12^ The TNBCtype4 classification system, for example, categorizes TNBC into four distinct subtypes: Basal-Like 1 (BL1), characterized by proliferative signatures; Basal-Like 2 (BL2), with growth-factor pathway activation; Luminal Androgen Receptor (LAR), associated with hormone-related genes; and Mesenchymal (MES), enriched for mesenchymal and epithelial-mesenchymal transition (EMT)-related signatures.^13^

Among these, the proliferative BL1 subtype is associated with favorable chemotherapy responses, while LAR subtypes show resistance^13^—paralleling the behavior seen in hormone receptor-positive breast cancers.^14–16^ While clinical use of these associations is currently restricted to HR+ HER2-cancers,^17^ a recent meta-analysis has demonstrated that nearly all breast-cancer mRNA-predictive-classifiers (TNBC and hormone positive), are primarily based on scoring estrogen- and proliferation-associated gene signatures.^18^

Despite these advancements, translating subtype classifications into clinical decision-making has been challenging in TNBC. Unlike hormone receptor-positive cancers, the higher cellular complexity of TNBC, which includes dense infiltrates of immune and stromal cells, complicates efforts to identify cancer-specific gene signatures. For example, the presence of immune cells and mesenchymal stromal components in tumor samples can obscure intrinsic subtype signals, as seen with the initial TNBCtype classification,^19^ which was refined to exclude immune-dominated and stromal-enriched subtypes to improve its clinical relevance.^13^

***Infiltrating Immune Cells*** are another layer of complexity in TNBC. Macrophages and fibroblast-derived interleukins, such as IL-6, have been implicated in promoting resistance to chemotherapy agents commonly used in TNBC treatment, including anthracyclines and platinum-based drugs.^20–22^ Interestingly, both anthracyclines and platinum agents can selectively target immune suppressive cells within the tumor, thus indirectly supporting anti-cancer immunity—a mechanism believed to underpin the success of some chemo-IO strategies.^23, 24^ In fact, both anthracyclines and platinum agents have been highlighted as potentially critical chemotherapy backbones underlying recently chemo-IO successes.^25^

Our hypothesis posits that both cancer-intrinsic subtypes and immune-cell composition of pre-treatment biopsy samples independently influence drug response in TNBC, underscoring the need for diagnostic approaches that evaluate both dimensions simultaneously. **CIBERSORT,** a widely adopted computational tool for immune cell quantification from bulk RNA-seq data, provides a pathway forward by deconvoluting immune cell composition within tumors.^26^ This approach has shown significant prognostic and predictive power in TNBC, highlighting lymphocytes and tumor associated macrophages (TAM) as critical components of the tumor microenvironment (TME).^27–29^

In our study, we validated that lymphocyte and TAM cell aggregates outperformed cancer-subtype signatures in predictive power. Additionally, we found that multicellular aggregate approaches that account for the spatial organization of the TME enhance reproducibility and accuracy across patient cohorts. By integrating spatial information from spatially resolved single-cell transcriptomics datasets, we reconcile past gene signature methods with contemporary spatial profiling technologies, offering a powerful approach to evaluating the spatial dynamics within the TME of bulk RNA-seq data and refining diagnostic approaches for TNBC. This holistic view on TNBC heterogeneity and immune dynamics not only provides a robust foundation for subtype-driven therapy selection but also paves the way for novel TME-modulating interventions aimed at improving therapeutic responses in this challenging cancer type.

## RESULTS & DISCUSSION

### Univariant Survival and Drug-response Analysis

To directly compare the predictive power of the CIBERSORT and TNBC-type classifiers, we curated datasets originally used to train the TNBC-type models and applied both the CIBERSORT and **TNBC-type** algorithms (Figure 1A, Figure S01), as outlined in the methods below. We generated aggregate tertiary lymphoid structure (TLS) and tumor associated macrophage (TAM) scores by averaging lymphocyte and M2-macrophage/monocyte scores, respectively. Univariate analyses of prognostic and predictive power were conducted using Cox and logistic regression models to derive hazard and odds ratios, respectively (Figures 1B and 1C).

**Figure 1.**
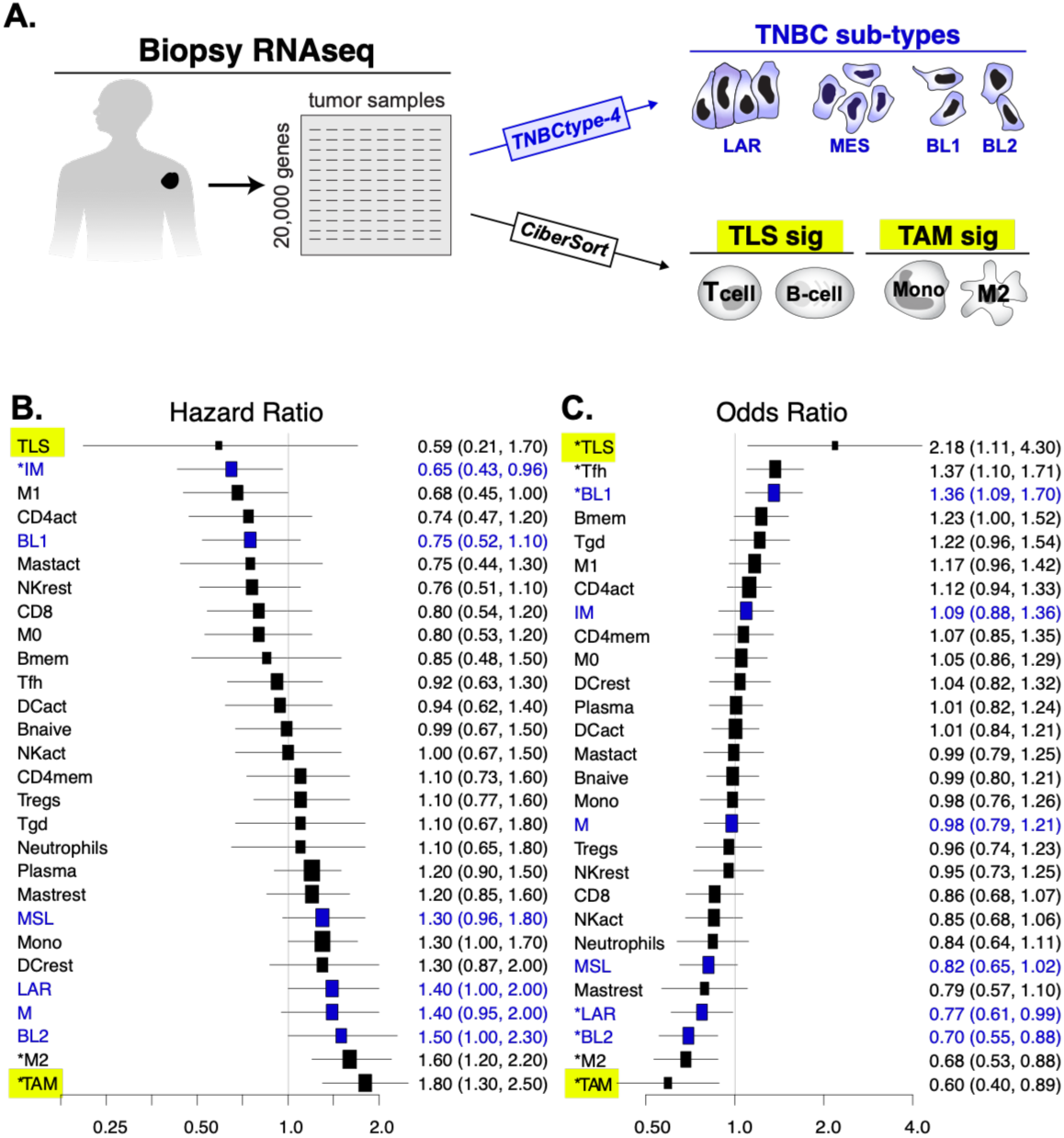
Overview of Prognostic and Predictive Power of TNBC-type and CIBERSORT signature-based methods. Statistically significant associations are marked with an *. **A.** Overall Strategy: two commonly used gene-signature based methods are evaluated on the same pre-treatment datasets **B.** Univariate survival analysis of CIBERSORT and TNBC-type signature enrichment across TNBC samples within TCGA breast cancer cohort. **C.** Univariate drug-response analysis of CIBERSORT and TNBC-type signature enrichment applied across 5 NAC-TNBC cohorts which were previously evaluated in the last refinement of the TNBC-type classifier.

Remarkably, TLS-like and TAM cell aggregates emerged as the strongest predictors of survival and drug response, exhibiting greater effect sizes and statistical significance in treatment response than the significant cancer subtype signatures (BL1, LAR, and BL2). Intriguingly, the Immunomodulatory (IM) TNBC subtype was uniquely associated with improved survival—a finding that underscores its relevance as a marker of TLS presence, even though it was omitted from the updated TNBC-type4 classifier due to its strong association with lymphocyte infiltration rather than cancer-intrinsic characteristics.^13^ This observation suggests that the IM subtype may serve as an effective TNBC-specific proxy for TLS-like structures.

While cancer subtype signatures demonstrated consistent predictive value, certain subtypes displayed distinct trends: the highly proliferative BL1 subtype was associated with favorable survival and drug response outcomes, whereas M/MSL/LAR/BL2 subtypes were linked to poorer responses. This pattern aligns with broader findings in breast cancer diagnostics, where high-proliferation and low-hormone gene expression profiles correlate with better outcomes in hormone receptor-negative cancers.^28, 29^

### Cell Mixtures Dynamics and Predictive Power

Our findings indicate that the predictive and prognostic power of gene signatures is predominantly driven by cell-type mixtures rather than pure cancer-type-specific signals. Tumors enriched with TLS-like cell aggregates (**blue**) tended to show improved survival and treatment response, whereas those dominated by TAMs (**red**) were associated with poorer outcomes (Figure 2A).

**Figure 2.**
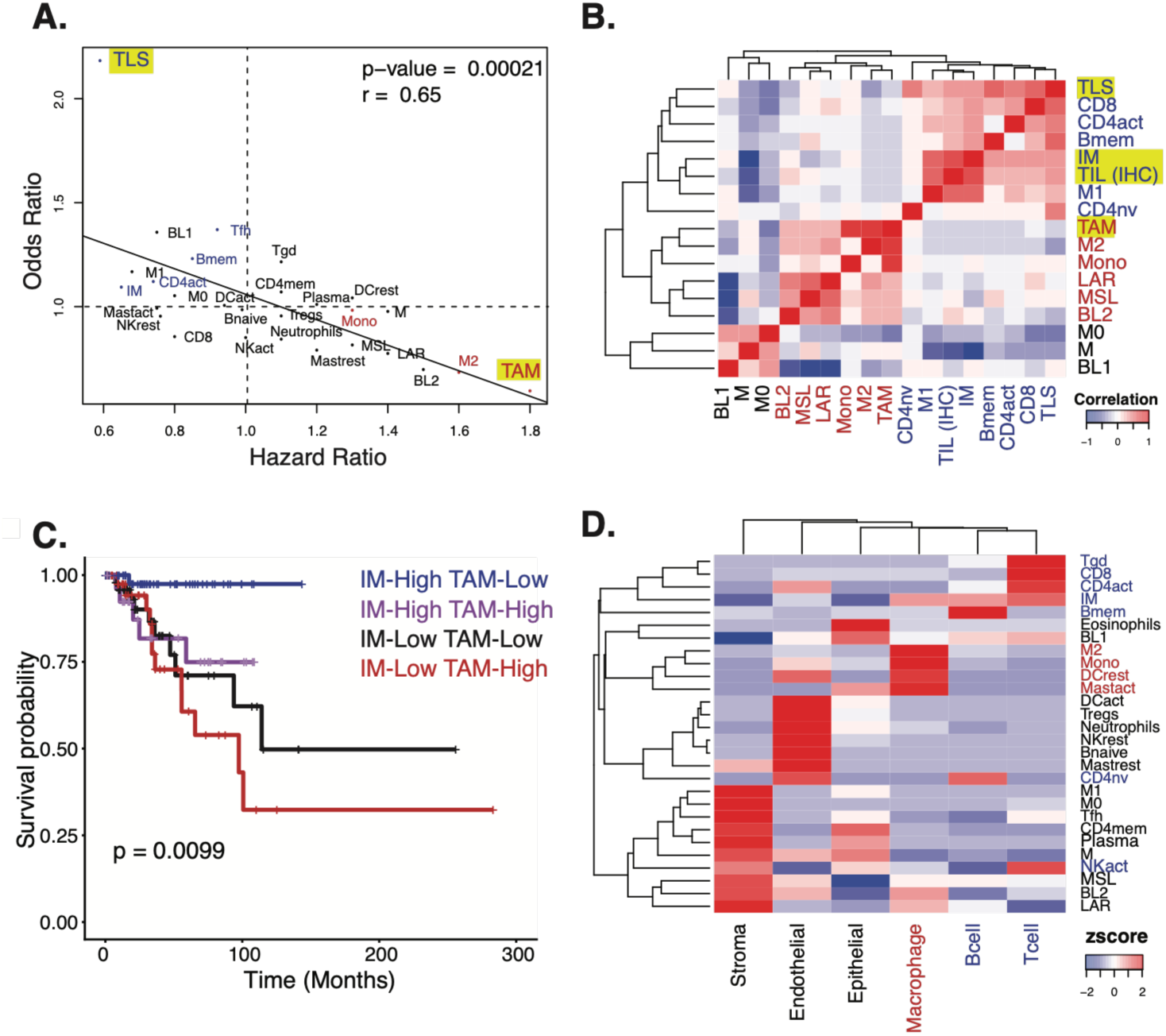
Performance of CIBERSORT and TNBC-type gene signatures largely driven by three sources of variance: Cancer, Macrophages and Lymphocyte signatures. **A.** Prognostic (survival) and predictive (drug-response) power of gene signatures is highly correlated across both CIBERSORT and TNBC-type **B.** Evaluation of TNBC subtype gene-signature and immune cell composition score correlation across patient cohorts reveals three clusters of predictive variance: Cancer, Macrophages and Lymphocytes. This analysis demonstrates that TAM and Lymphocyte content are major sources of independent variability. **C.** Kaplan-Meier visualization of independent predictive power of TAM and IM quantified using Cox-regression. **D.** Enrichment scores (z-score) TNBC-type signatures and immune cell composition scores indicate that predictive power of each component may be driven by mixtures of cells.

Notably, combined TLS and TAM scores enhanced predictive power relative to individual immune cell signatures, underscoring the complex cellular interactions within the TME. This correlated predictive and prognostic power was highly statistically significant (r = 0.65, p-value = 0.00021) and closely mirrored that of OncotypeDx in hormone-positive breast cancer, suggesting that similar underlying principles may apply across distinct cancer contexts.

**IHC-defined TILS:** To explore the structure of these gene signatures further, we examined their correlations within the TCGA cohort, which also included immunohistochemical quantification of TILs (Figure 2B). This analysis revealed three main clusters based on TLS-like scores, TAM content, and cancer subtype. Interestingly, the IM gene signature displayed the strongest correlation with IHC-TIL, affirming its role as a marker for TIL levels in TNBC. The relatively weaker correlation of TAM signatures with BL2, MSL, and LAR subtypes may reflect the presence of macrophage and stromal gene contributions within these subtype profiles (Figure 2B).

Given the high predictive power of the IM subtype and TLS-like and TAM aggregates in our univariate analysis (Figure 1 B-C) and their largely independent effects (Figure 2B), we evaluated their combined prognostic utility using multivariate Cox regression. The combination of TAM aggregates and IM subtype yielded the strongest predictive model (p-value = 0.0099), driven by independent contributions from both TAM (p < 0.001) and IM (p = 0.048) signatures. Kaplan-Meier survival curves further illustrated the robust predictive capacity of these signatures, with stratification based on high and low TAM and IM scores (Figure 2C).

### Impact of TME Complexity on Signature Interpretation

We next investigated the effects of stromal and tumor-infiltrating immune cells on CIBERSORT and TNBC-type classification using scRNAseq obtained from TNBC patients (Figure 2D, S2-3).^30, 31^ Cell composition scores and gene signature scores derived from both CIBERSORT and TNBC-type mapped across epithelial, macrophage, stromal, and lymphocyte cells, revealed that most “pure cell” signatures align with mixtures of cell types rather than pure populations (Figure 2D). For instance, the IM signature, strongly associated with immune infiltrate, correlated with combined signatures from T-cells, B-cells, and macrophages. This supports the idea that these gene signatures capture complex TME niches rather than isolated cell types. The sole exception was the CD8 and M2 macrophage signatures which exhibited a clear association with pure populations in TNBC, underscoring their unique role within the TME.

Our analysis revealed significant enrichment of stromal cells in several TNBC-type subtypes (MSL, BL2, LAR, and M) and CIBERSORT immune cell scores, including M1/M0 macrophages, CD4 memory cells, and T-follicular helper cells. This enrichment suggests that stromal infiltration may confound both cancer and immune classifications due to overlapping gene expression across these compartments. To visualize this gene expression overlap, we examined compartment-specific gene expression across 20,000 protein-coding genes, as shown in Figure S2A-B. Interestingly, stromal cells demonstrated the greatest gene expression overlap with the epithelial compartment, highlighting the challenge of distinguishing stromal and epithelial contributions in TNBC classification.

To further explore the influence of stromal and immune genes on TNBC-type classifications, we systematically removed high-expressing genes from stromal, lymphocyte, and macrophage compartments (Figure S2C). Remarkably, the removal of T-cell, stromal, and macrophage genes had a more substantial impact on the fidelity of TNBC-type classification than the removal of cancer-specific genes alone, emphasizing the influence of the tumor microenvironment (TME) on TNBC subtype fidelity. Additionally, TNBC-type stromal genes appeared largely independent of hormone and interferon-related pathways, which are primarily enriched in LAR cancers and lymphocyte populations, respectively(Figure S2D). These findings suggest that inflammatory and mesenchymal gene sets significantly contribute to the predictive power of TNBC-type signatures (Figure S2E), supporting the hypothesis that bulk RNA-seq classifiers capture the broader “ecosystem” surrounding cancer subtypes.

This observation argues for the utility of the TNBCtype-IM signature which was removed in TNBC-type4.

### Validation of Retrospective Results using in-house RNAseq and IHC

To validate our retrospective reconciliation of gene-signature and immunohistochemistry (IHC) results, we used an in-house dataset of 25 TNBC samples treated with the ACT (Doxorubicin, Cyclophosphamide, Paclitaxel) regimen. This dataset included paired pre-treatment biopsies and surgical samples, which were sequenced and assessed with IHC. We expanded our validation by collecting demographic, treatment, and response data for a total of 67 TNBC patients. This dataset included patient age, diagnosis date, BMI, cancer grade, stage, and treatment response data, as detailed in Supplementary Table 1. Most patients received anthracycline- and taxane-based neoadjuvant chemotherapy (NAC), with ACT being the most common regimen. Post-treatment, 27 patients (40%) achieved pathological complete response (pCR), while 40 (60%) had residual disease. RNA was extracted and sequenced from 42 pre-treatment biopsies and 31 residual samples from these patients. Overall, TLS-like and TAM aggregate odds ratios in this cohort showed rankings consistent with our retrospective analysis, although only the IM signature achieved statistical significance (Figure 3A-B).

**Figure 3.**
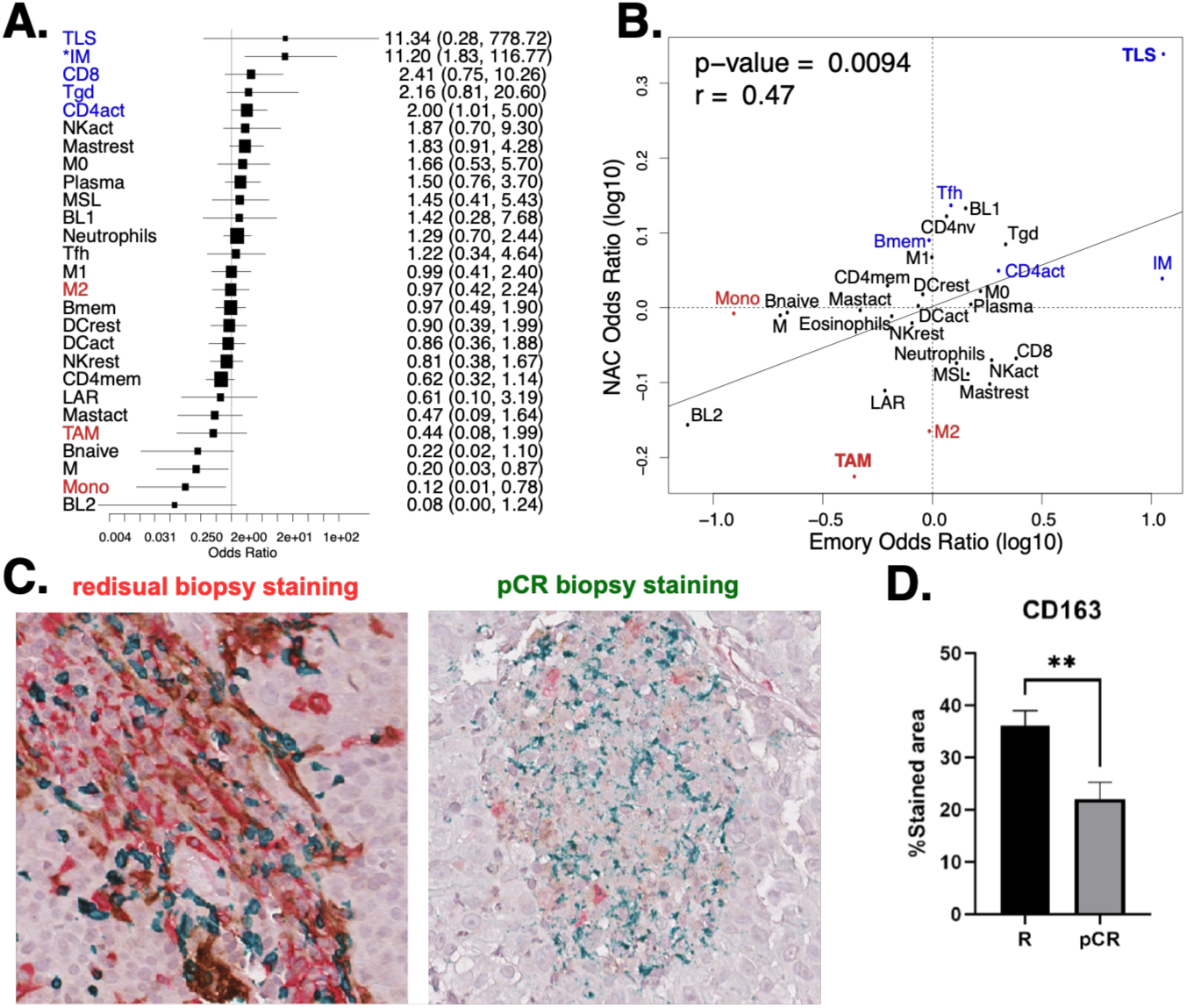
Validation of TLS and MDSC associations with new patient cohort treated with ACT. **A.** Evaluation of univariate predictive power of CIBERSORT and TNBC-type showed similar rankings of TLS-like, TAM and Cancer subtypes even though only the IM subtype showed the only statistically significant association. **B.** Signature rankings between Emory and NAC cohorts were highly correlated despite variability of individual signatures. **C-D.** IHC analysis on the new patient cohort validates that M2-macrophage infiltrate correlates with residual disease following ACT treatment.

We also evaluated pre-NAC (ACT) biopsy specimens for M2 macrophages, cytotoxic T cells, and PD-L1 expression using multi-color multiplex IHC with CD163 (clone SP57), CD8 (clone MRQ26), and PD-L1 (clone SP263) antibodies. Freshly cut sections from these biopsies underwent color-based k-means segmentation to quantify antibody staining across the tumor section. Multivariate regression analysis indicated that TAM markers (CD163) were significantly associated with residual disease (p = 0.0392), while TIL-associated markers (CD8, p = 0.2669; PD-L1, p = 0.8437) were not significant. Representative images with varying CD163 (red), CD8 (green), and PD-L1 (brown) expression patterns are shown in Figure 3C, with total CD163 quantification and variability depicted in Figure 3D.

### Validation of TLS-,TAM- and cancer-stromal niches within the TME (**Figure 4**)

Overall, a paradoxical finding of this study was that while TLS- and TAM- and cancer-subtype showed statistically significant predictive and prognostic power, genes from other cell-compartments appeared to be important to the predictive power of these signatures. This led us to hypothesize that the tumor-microenvironmental structure itself might reflect the cell-type makeup of our predictive signatures.

**Figure 4.**
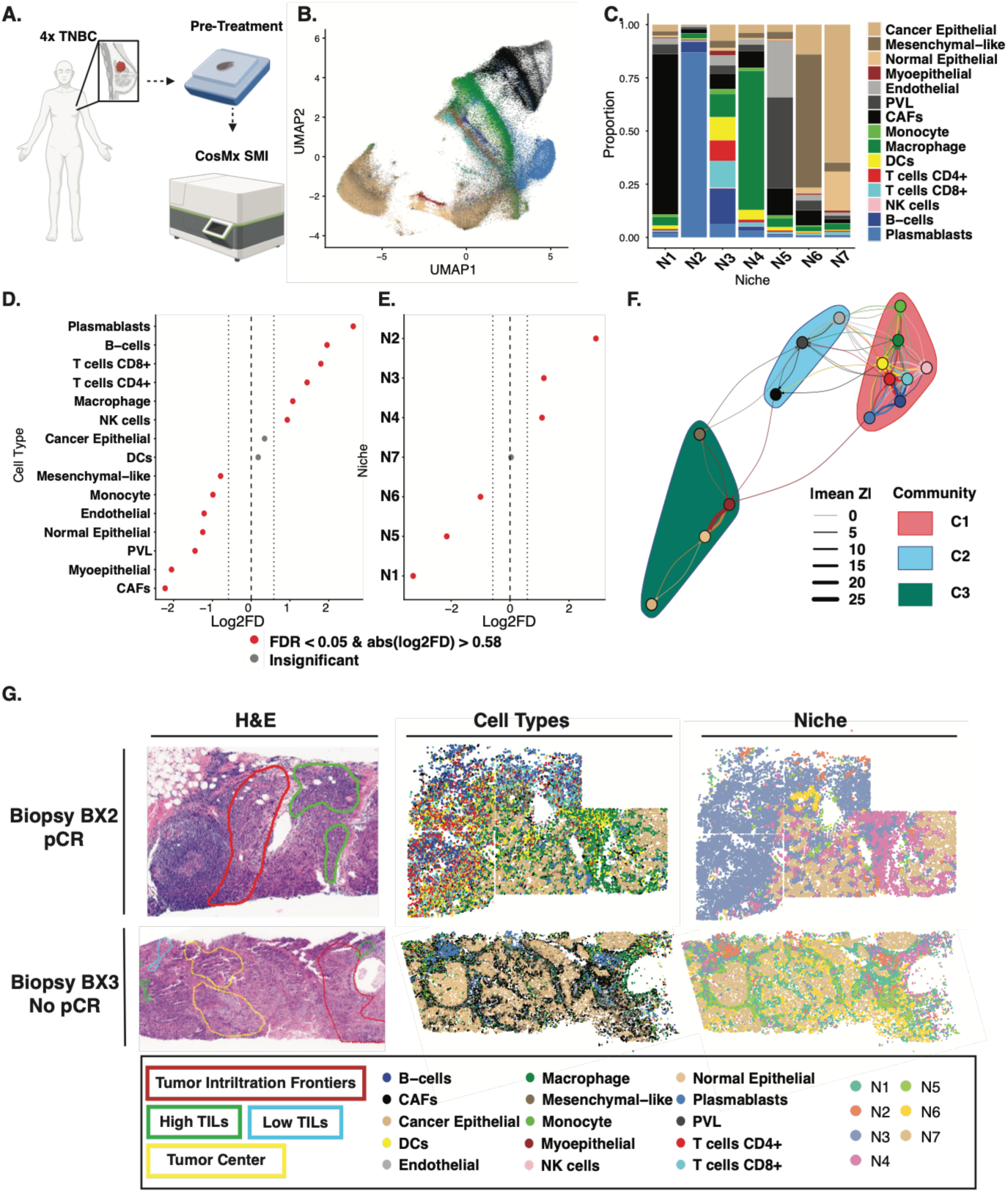
Validation of TLS-, TAM- and Tumor-stromal niches within the TME. **A.** Schematic representing four TNBC pre-treatment biopsy sample collection and profiling using CosMx Spatial Molecular Imager. **B.** UMAP projection of 15 unique cell types identified across 4 TNBC patient samples. **C.** Cell composition of 7 niches identified across all 4 samples. **D.** Permutation-based testing reveals strong lymphocyte associations with pCR while stromal cells and monocytes have significant association with no pCR. **E.** Permutation-based testing reveals strong associations of immune-rich niches (niches 2,3,4) with pCR while stromal and mesenchymal-like epithelial niches (niches 1,5,6) have strong associations with no pCR. **F.** Consensus colocalization networks generated from cross-PCF analysis reveal strong spatial enrichment of immune cells and separate colocalization of epithelial cells and stromal cells across all samples. Cells/nodes resolve into communities with strong internal colocalization trends and variable external colocalization trends. **G.** H&E, cell types, and niches of representative pCR (top row) and no pCR (bottom row) biopsy samples highlighting the contrast between immune cell and immune niche abundance in the tumor microenvironment.

To test this hypothesis, we examined the spatial organization of the TNBC microenvironment using subcellular resolution spatial transcriptomics (CosMx Spatial Molecular Imaging (SMI)) with the 1k gene panel. Here, 4 pre-treatment TNBC biopsy samples were selected where 2 of which were obtained from patients that would achieve a pathological complete response (pCR) after treatment and two were non-responders (No pCR) to treatment. Patients were treated with a combination of chemotherapy and anti-PD1 immune checkpoint blockage in the form of pembrolizumab. All 4 samples were profiled using the CosMx spatial molecular imager yielding over 350,000 cells across 500 FOVs. 15 major cell types were identified including tumor epithelial cells, key lymphocytes found in TLS-like structures, phenotypically plastic myeloid cells, and stromal cells(Figure 4A-B). Permutation testing of cell type proportions in pCR vs No pCR samples revealed strong lymphocyte preference for pCR samples, while stromal cells such as cancer associated fibroblasts (CAFs), perivascular cells, and endothelial cells were highly enriched in No pCR samples (Figure 4D). Interestingly, macrophages are more associated with pCR while monocytes were more associated with No pCR. While M1 and M2 phenotypes were not distinguished in this cohort, differentially expressed macrophage genes included both proinflammatory/M1 (NR1H3+) and pro-resolution/M2 (SPP1+, APOC1+) gene expression patterns, indicating a continuous M1/M2 functional spectrum (Figure S06 A).

#### Spatial Aggregate Characterization Methods

We next characterized cellular aggregation within these samples using two unbiased, complementary methods: (1) niche analysis and (2) spatial point-statistical analyses. Niche-based analysis involves clustering of cellular neighborhoods into larger regions of common colocalization patterns which reveals broad tissue structures and functional immune niches. Spatial statistical analysis, specifically the cross-pair correlation function, offers a complementary approach to quantifying colocalization by measuring colocalization of cell populations across different length scales.

While niche-based analysis is common practice in spatial transcriptomic analyses, number and composition of niches heavily depend on user-defined parameters (neighbor distance and cluster number), which dramatically affect the scale of detected tissue structure. Spatial summary statistics are an emerging alternative that quantify colocalization across continuous ranges, avoiding the need to pre-specify neighborhood scale.

These methods capture different features of spatial organization. Niches represent small recurring regions at a user-defined scale, while cross -pair correlation summarizes co-localization across a range of distances, not just within user-defined neighborhoods. Furthermore, networks resolve into communities or clusters where highly colocalized cell populations engaged in common biological processes. We apply both methods to identify colocalization trends that support the presence of multicellular aggregates identified in bulk samples.

### Spatial Niche Analysis

Multicellular niches across all samples were identified by clustering individual cell neighborhoods. Cellular neighborhoods were determined by identifying cells found within a 15 µm range of a given cell. K-means clustering across all four samples revealed 7 spatially and functionally independent niches (Figure 4 C-G, S05 C-D). Among these niches were two epithelial-dense niches (N6-7), three immune niches (N2-4), and two stromal niches representing vascular structures (N5) and peripheral stromal structures (N1).

Among the immune niches, N2 was composed primarily of plasmablasts found in the peripheral stroma and was most abundant in pCR samples (Figure 4E, S05 E). Niche N3 was composed primarily of T- and B-lymphocytes, antigen presenting cells, and structural stromal cells, resembling TLS-like cell aggregates. Permutation testing revealed that niche N3, though present in all samples, was significantly more abundant in pCR samples, consistent with the relationship between TLS-like cell aggregates and treatment response characterized in Bulk RNA-seq analysis (Figure 4E). Niche N4 was dominated by macrophages and functionally heterogeneous with significant enrichment of Type I interferon response, inflammatory response, and angiogenesis gene sets among other hallmark pathways (Figure S05 C). Niche N4 was also highly associated with pCR samples, possibly reflects the paradoxical function of M1 and M2 macrophages in the TME.

### Spatial Statistical Analysis

To complement the niches identified through neighborhood clustering, we applied the cross-pair correlation function (cross-pcf) between all cell populations to generate colocalization networks. We computed the mean cross-pcf for all cell type pairs across a 50µm range for each sample individually. Sample-specific colocalization networks (Figure S06 F-I) and consensus networks (Figure 4 F, Figure S06 A,C) reveal consistent and strong colocalization within immune cells, stromal cells, and epithelial compartments. Notably, CD8+ T cells, CD4+ T cells, B-cells, and dendritic cells (DCs) were among the most influential nodes, with strong colocalization patterns between themselves and with macrophages and monocytes (Figure S06 A,B). Node diversity revealed that myeloid cells such as DCs, macrophages, and monocytes were often colocalized with several different cell types relative to other cell populations (Figure S06 A,B). This may reflect the phenotypic heterogeneity of TAMs and myeloid cells in general in the tumor microenvironment. Critically, both niche-analysis and these colocalization patterns confirm the spatial patterns we speculate are captured in TLS-like and TAM aggregate signatures in our bulk analysis.

We then applied the infomap community detection algorithm to find groups of densely connected nodes, revealing cell populations that tend to colocalize with each other. Remarkably, networks resolve into communities that reflect the spatial niches identified through clustering cell neighborhoods. Three communities were identified in the consensus colocalization network where immune cells were organized into the most centralized and cohesive community, C1, while communities C2 and C3 were primarily composed of stromal and epithelial cells, respectively (Figure S06 C-E). Nodes within communities C1 and C3 preferentially colocalize with themselves rather than nodes in other communities, indicating strong immune/tumor compartmentalization in the tumor microenvironment (Figure S06 D). In contrast, nodes in community C2 display comparable levels of colocalization with nodes within their own community and nodes in other communities. Together with the high polarity of community C2, this pattern indicates that stromal cells often serve as the interface between immune and tumor compartments in the tumor microenvironment. Communities in individual samples were consistent in immune and epithelial composition, with slight variation in stromal community composition across samples, generally involving reassignment of CAFs to tumor and immune dominant communities (Figure S06 F-I).

## CONCLUSION

In conclusion, the challenge of precisely matching cytotoxic chemotherapies to TNBC patients remains substantial, especially given the shift to a standard-of-care treatment combining chemotherapy and immunotherapy for this patient population. While both cancer subtype and infiltrating immune cells have been demonstrated to correlate with survival and treatment response, neither alone has achieved sufficient predictive power to serve as a reliable diagnostic tool. We hypothesize that both infiltrating immune cells and cancer subtype contribute independently to treatment outcomes, necessitating diagnostics based on gene signatures that account for both factors.

Our study reinforces this hypothesis by showing that both cancer subtype and immune cell infiltration can predict response to therapy; however, immune infiltrate—particularly macrophages and lymphocytes—play a larger role. Through detailed gene signature analysis, we identified that tumor niches defined by multi-cellular immune and cancer cell interactions show improved reproducibility and predictive power across patient cohorts. Specifically, these niche-based signatures revealed robust contributions from both immune cell clusters and distinct tumor environments, with macrophages and lymphocytes showing independent, predictive influences on treatment outcomes.

This finding supports the view that multi-cellular spatial organization within the TME holds significant prognostic value, reconciling discrepancies observed in earlier gene-signature methods and offering a new framework for evaluating spatial logic in bulk gene signatures. Given that both cancer and immune cell phenotypes are intrinsically multi-cellular and highly contextual, our niche-based approach aligns with the biological reality of the TME as a complex ecosystem.

These insights open pathways for designing interventions that modulate the TME to enhance anti-cancer efficacy. By targeting specific immune cell types, such as macrophages and lymphocytes, within their spatial niches, future therapies may improve responses in TNBC and potentially other cancers by leveraging the multi-cellular dynamics of the TME. Our study’s framework offers a platform for future work aiming to characterize and manipulate the TME to achieve more tailored and effective anti-cancer strategies.

## MATERIALS AND METHODS

### Patient, specimens, and response assessment

This study was approved by institutional review board of Emory University and The Ohio State University. 67 triple-negative invasive breast carcinoma patients treated with neoadjuvant chemotherapy and follow-up surgical resection (39 lumpectomy specimens and 27 mastectomy specimens) were included in this study. For neoadjuvant chemotherapy, all patients received four cycles of AC (doxorubicin+cyclophosphamide) followed by paclitaxel. Surgical resection specimens were evaluated, and pathologic complete response (pCR) was defined as no residual invasive carcinoma in the breast and no lymph node metastasis.

### Tumor RNA extraction and sequencing

Tumor RNA extraction and QC were performed at the Emory Integrated Genomics Core (EIGC) utilizing the E.Z.N.A. FFPE RNA kit (Omega Biotek). RNA sequencing was performed at Discovery Life Sciences. The concentration and integrity of the extracted total RNA were estimated using the Quant-iT™ RiboGreen® RNA Assay Kit (Thermo Fisher Scientific) and the Fragment Analyzer Systems using the RNA High Sense chip, respectively. Up to 30ng of RNA with DV200>20% was used to prepare libraries using SMRTer Stranded Total RNAseq v2 (Clonetech) with slight modifications. Post-ligated material was individually barcoded with unique dual indexed primers. The concentration of the libraries was then measured by Picogreen assay (Thermo), and the average fragment size of the libraries was estimated by utilizing a DNA High Sense chip on a LabChip GX Touch Nucleic acid analyzer (PerkinElmer), respectively. KAPA qPCR assay (Roche) was performed to assess the nanomolar amounts of ligated libraries. Final libraries were pooled equimolar and sequenced as 2×100bp Paired-end sequencing on the NovaSeq 6000 instrument using an S4 200 cycle flow cell to achieve the desired read depth.

### Immunohistochemistry

IHC was performed using biopsy tissues with three antibodies including CD163 (clone SP57, rabbit; Ventana Medical Systems, Inc), CD8 (clone MRQ26, mouse; Ventana Medical Systems, Inc,) and PD-L1 (clone SP263, rabbit; Ventana Medical Systems, Inc, Tucson, AZ, USA). M2 macrophages were identified by positive CD163 staining.^32, 33^ We evaluated M2 macrophages, cytotoxic T cells and PL-L1 expression by applying a multi-color multiplex IHC. M2 macrophages and cytotoxic T cells were evaluated by estimating the percentage of tumoral or stromal areas infiltrated by CD163-positive cells and CD8-positive cells, respectively. PD-L1 expression was evaluated by estimating the percentage of tumor cells or stromal (immune) cells with membranous PD-L1 staining. Representative images with different combinations of CD163 (red), CD8 (green) and PD-L1 (brown) expression are illustrated in Fig. 3C. Statistical analysis was performed using SAS version 9.4 for Windows (SAS Institute, Inc, Cary, NC, USA). Descriptive statistics were used to summarize patient clinical and pathologic characteristics. Categorical data were summarized as frequency and percentage, and continuous variables as medians and ranges. To study the associations with pCR, Fisher’s exact test was used for categorical variables. Wilcoxon rank-sum test was used to compare the continuous variables. All the continuous variables have been tested for normality using Kolmogorov–Smirnov test. A multivariable logistic regression model was used to determine the variables associated with the incidence of death as well as recurrence. Variables with a *p* value < 0.10 in the univariate analysis were entered into a multivariable model. Variables were removed sequentially from the multivariable model using the backward selection method.

### Color-based K-means Segmentation

Firstly, at most 10 image patches with 512×512 pixels in size were selected from each IHC tissue with the lowest excluded region ratio. Secondly, we converted all selected image patches from RGB color space to L*a*b* color space, which ensures the highest color contrast across three different IHC markers. Thirdly, k-means clustering was performed and aggregates each pixel of selected patches in L*a*b* color space. In detail, we set K = 15, number of initializations = 3, and maximum number of iterations = 300 with tolerance = 1×10^−4^. Next, two pathologists helped identify and confirm clustering centers for CD8, CD163, and PD-L1. Each IHC markers may contain several clustering centers. Finally, given the learned k-means clustering center, we applied the rest of the IHC WSIs to this model and obtained the IHC markers segmentation results.

### Statistical Analyses

#### Data Availability Statement

The data generated in this study are available within the article and in the supplementary data files. Raw data is also available through the Gene Expression Omnibus (https://www.ncbi.nlm.nih.gov/geo/) under accession number GSE260693. CosMx spatial transcriptomics data is available upon request.

#### Data and Code Availability

Data visualization and analysis code is publicly available through a GitHub Repository (https://github.com/wcb96858/TNBC_Analysis). All data preparation procedures, data analysis procedures, and visualization were conducted using Rstudio^34^ and are described, in detail, below. Corresponding R scripts are stored in the GitHub Repository as mentioned before.

#### Gene Expression, Survival and Drug-Response Datasets

This study used data from the public domain that had been previously used to evaluate the most recent version of TNBC-type algorithm TNBC-type4.^13^ Use of this data was critical to benchmark the performance of intrinsic (TNBC-type4) and extrinsic (CIBERSORT) classification methods in this study. TCGA gene expression and survival data were obtained for Breast Invasive Carcinoma (TCGA, PanCancer Atlas) from http://www.cbioportal.org/datasets.^35, 36^ RSEM mRNA expression data and clinical survival data were used for classification and prognostic analyses described below. A subset of 179 triple-negative breast cancer (TNBC) samples defined in the latest TNBC-type4 publication^13^ were the focus of all analyses. TCGA data for these 179 samples were sliced out and stored with previous subtype-scores. Pre-Neoadjuvant Chemotherapy (NAC) treated mRNA expression with paired NAC-response data was obtained as previously described in the most recent TNBC-type4 publication.^13^ Briefly raw microarray data, was obtained from Gene Expression Omnibus (https://www.ncbi.nlm.nih.gov/geo/) under accession numbers GSE25066,^37^ GSE41998,^38^ GSE22358,^39^ GSE22226^40^ and GSE32646.^41^ Hugo gene-symbols were mapped to probe ids, and genes with multiple ids were averaged to yield consensus gene expression for each unique gene. NAC response data was stored as a named vector where complete pathological response (pCR) was encoded as “1” and residual disease was encoded as “0”. TNBC-patients within each cohort were defined based on definitions published in the most recent TNBC-classification paper.^13^ Gene expression and NAC-response for each dataset were saved as RData files named by their GEO accession numbers. Reads for in-house RNA-seq validation cohort of 42 pre-treatment biopsies and 31 resected tumors were pseudo aligned using Kallisto.^42^ *Single-cell RNAseq Breast Cancer Data* was obtained from GEO under accession numbers GSE118389^43^ and GSE176078^31^. Published cell-type annotations were mapped to sample id’s and “pseudo-bulk” matrices were constructed for each cell type by average TPM values across each annotated cell-type.

#### CIBERSORT and TNBC-type classification procedures

CIBERSORT deconvolution of bulk samples and TNBC-type gene-signature enrichment scoring was determined based on previously published procedures. *CIBERSORT source code* was obtained by request from https://cibersortx.stanford.edu/contact.php and modified slightly to add a progress bar for very long calculations.^26^ Previously published cancer (LM6) and immune-cell (LM22) signature matrices were obtained from the CIBERSORT website. Two breast cancer cell type signatures were generated using the signature-generation tool hosted on the CIBERSORT website. These signatures were derived from breast cancer scRNA-seq datasets: GSE118389^43^ and GSE176078^31^. The CIBERSORT algorithm was applied to all NAC-datasets, using all 4 signatures described above. *TNBC-type signature enrichment and classification* was performed using a modified procedure published by Karaayavaz and coworkers.^43^ Briefly, gene-lists, direction, and Bonferroni-corrected p-values were extracted from Chen and co-workers^19^ to obtain complete gene-signature information for BL1 (678 genes), BL2 (439 genes), IM (1067 genes), LAR (2236 genes), M (856 genes), MSL (2365 genes). This compiled data was used to generate networks for single-sample enrichment analysis using the VIPER algorithm.^44^ Briefly, the Viper regulon objects for each signature were defined by assigning ‘tfmode’ based on log2FC sign (+1 or -1) and ‘likelihood’ was defined as (1 - p-value) of differential expression. TNBC-signature enrichment was calculated for all NAC-datasets. *Combined TNBC-type and CIBERSORT scored matrices* were normalized and concatenated with treatment-response vectors to enable univariate and multivariate linear and logistic regression as described below.

#### Prognostic and Predictive Analysis

Comparison of the prognostics and predictive power of the CIBERSORT and TNBCtype signature enrichment scores was conducted on TCGA and NAC datasets outlined above. These datasets were used to evaluate prognostic and predictive power of the most recent revision of TNBCtype^13^ and are important to benchmark the performance of both gene-signature based classifiers. *TCGA Prognostic power of signatures* was evaluated using 179 TNBC patient id’s defined in previous TNBCtype publication.^13^ Briefly, TCGA patient clinical data was modified to include previously published TNBC-type signature scores, TLS-IHC scores^13^ and CIBERSORT scores using the LM22 immune-cell signature matrix. Univariate COX regression was applied across all TNBCtype and CIBERSORT signatures, and p-values and hazard ratios were stored in matrix form to enable visualization (Figure 1B-C). Univariate and multivariate survival analyses and visualization were conducted using Cox regression and Kaplan Meier curves from the rms^45^ and survminer^46^ R packages. Clustering of signature correlations was visualized using the gplots^47^ and RColorBrewer^48^ packages. *NAC-treatment Predictive power of signatures* was evaluated using the 5 NAC datasets defined above. Signature scores paired with treatment response were used to evaluate the univariate predictive power of all signatures via logistic regression using the glm function within base R.^49^ Odds ratios and p-values were calculated for each signature within each NAC dataset. Consensus odds ratios and p-values were obtained across all datasets through Stouffer-integration. Final consensus values were stored in matrix form for visualization. Correlation of odds and hazard ratios were conducted using the ‘lm()’ function in base R.

### scRNAseq Analysis

Analysis of TNBC-type and CIBERSORT signature enrichment on scRNAseq datasets was conducted with the two breast cancer datasets described above. ^31, 43^ Briefly, TNBC-type and CIBERSORT enrichment scores were calculated on both datasets using methods described above. Signature scores were concatenated on overlapping cells and raw enrichment scores normalized by taking the z-score of each row. Normalized Enrichment scores were visualized using the gplots package (Figure S02D). Venn Diagrams of Cell-type specific genes were generated using the eulerr^50^ package in R. For each assigned cell type, the total expression of each gene within a cell type was divided by the sum of the gene-expression across all cell types. Cell-type specificity for cancer, macrophage, T-cell, B-cell, fibroblast, and endothelial compartments were defined by setting a threshold of 0.3 where total gene expression greater than 30% associated with one compartment was assigned to that compartment. This enabled visualization of all genes where a plurality of the expression was restricted to one or two cell-types (Figure S02B, E)

#### Cell-type specific gene-pruning of TNBC-type signatures

TNBC subtype classification fidelity was evaluated through removal of genes specific to six cell types: macrophages, stromal cells, T-cells, B-cells, endothelial cells, and epithelial cells. Here, a pseudo bulk dataset was derived from averaging normalized counts for each gene within cell types in a single-cell RNA-seq matrix. For each cell type, genes in the pseudo bulk set were ordered based on descending normalized counts and iteratively removed from TNBC subtype genes sets, followed by single sample enrichment analysis on 179 TNBC samples from TCGA using R package, Viper.^44^ Upper and lower bounds were determined by removing TNBC subtype specific genes from gene sets based on decreasing and increasing p-values, respectively. Subtypes were assigned to samples based on highest enrichment across the four subtypes. Fidelity scores were defined by the total change in number of samples classified by each subtype for each additionally removed gene with respect to initial classifications of samples. Fidelity scores were calculated for removal of cell type specific genes as well as TNBC subtype specific genes. Changes in fidelity score as the number of stromal, macrophage, T-cell, and epithelial (cancer) genes removed increases were visualized with respect to pruning of TNBC-specific genes. (Figure S02 C).

#### Pathway Analysis of TNBC-type and CIBERSORT gene-signatures

Pathway analysis of TNBC subtypes signatures were conducted on TNBC subtype-labelled samples from TCGA. Labels were determined by, first, reducing TNBC subtype gene signatures to significant differentially expressed genes (p-value < 0.05, n = 1582 total genes) and running VIPER on the 179 TNBC samples. Subsequently, the subtype with the highest enrichment for each sample was assigned as the corresponding label. Average TPMs across samples of each subtype were condensed into an expression matrix with each column representing a subtype where rows were the average expression of genes for a given subtype. This average subtype expression matrix was combined with the pseudo bulk expression matrix of six cell types curated during pruning, followed by VIPER analysis of standard hallmark pathway gene sets from the Molecular Signatures Database (MSigDB).^51, 52^ Scaled pathway enrichment was visualized and clustered using the gplots^47^ R package (Figure 3D).

### Spatial Sample Processing

#### Sample preparation for CosMx™ SMI RNA assay

To prepare the tissue samples for the CosMx™ SMI platform (NanoString Technologies Inc, Seattle, WA), samples were baked overnight at 60°C to ensure formalin-fixed, paraffin-embedded (FFPE) tissue core adherence to the glass slides. Samples underwent deparaffinization, proteinase K digestion, and heat-induced epitope retrieval (HIER) procedures to expose target RNAs and epitopes using the Leica Bond Rx system. For breast cancer samples, 1 μg/ml of proteinase K (ThermoFisher) was incubated at 37°C for 15 min and HIER at 100°C for 10 min in Leica buffer ER1 conditions were used. The samples were rinsed with Phosphate-buffered saline (PBS) once before incubating in 1:1000 diluted fiducials (Bangs Laboratory) in 2X SSCT (2X saline sodium citrate, 0.001% Tween 20) solution for 5 min at room temperature. Excessive fiducials were removed by rinsing the samples with 1X phosphate-buffered saline (PBS), followed by fixation with 10% neutral buffered formalin (NBF) for 1 min at room temperature. Fixed samples were rinsed with Tris-glycine buffer (0.1M glycine, 0.1M Tris-base in DEPC H2O) and 1X PBS for 5 min each before being blocked using 100 mM N-succinimidyl acetate (NHS-acetate, ThermoFisher) in NHS-acetate buffer (0.1M NaP, 0.1% Tween PH 8 in DEPC H2O) for 15 min at room temperature. Prepared samples were rinsed with 2X saline sodium citrate (SSC) for 5 min, and then an Adhesive SecureSeal Hybridization Chamber (Grace Bio-Labs) was placed to cover the samples.

1000-plex RNA ISH probes were denatured at 95°C for 2 min and then placed on ice before preparing ISH probe mix (1 nM ISH probes, 1X Buffer R, 0.1 U/μL SUPERaseIn™ in DEPC H2O). The ISH probe mix was pipetted well and applied to the tissue, covered with an Incubation Frame Cover to prevent evaporation. Then, the slides were placed in the hybridization chamber at 37°C for overnight incubation in the hybridization oven. After the overnight hybridization, samples were washed with 50% formamide (VWR) in 2X SSC at 37°C for 25 min twice and rinsed with 2X SSC for 2 min twice at room temperature. After blocking, the samples were washed twice using 2X SSC for 2 min at room temperature. A custom-made slide cover was attached to the sample slide to form a flow cell.

### SMI instrument run

The 980-plex RNA assay was designed to detect 960 target genes and 20 negative probe controls. Negative control probes are modeled after synthetic sequences from the External RNA Controls Consortium (ERCC)^53^. RNA target readout on the SMI instrument was performed following published protocols^54^. In brief, samples in the assembled flow cell were loaded onto the SMI instrument and washed with a Reporter Wash Buffer to rinse the samples and remove air bubbles. The entire flow cell was scanned (SMI preview scan) for RNA readout. The RNA readout cycle started by flowing 100 μl of Reporter Pool 1 into the flow cell and incubating for 15 min. After incubation, 1 mL of Reporter Wash Buffer was flowed into the flow cell to wash out the unbound reporter probes, followed by replacing the Reporter Wash buffer with Imaging Buffer prior to imaging. Nine Z-stack images (0.8 μm step size) of each FOV were acquired, and then fluorophores on the barcoded reporter probes were released at photocleavable linkers by UV illumination and washed off with Strip Wash buffer. This fluidic and imaging procedure was repeated for the 16 reporter pools, and the 16-round reporter hybridization-imaging cycle was repeated multiple times to increase RNA detection sensitivity.

After RNA readout, samples were incubated with a 4-fluorophore-conjugated antibody cocktail against CD298/B2M, PanCK, and CD45 proteins and DAPI stain in the SMI instrument for 1 hr. Nine Z-stack images for 4 channels were captured after unbound antibodies and DAPI stain were washed with Reporter Washing Buffer, and the flow cell was filled with Imaging Buffer.

#### <u>CosMx Data Analysis</u> Initial Data Processing

Four TNBC pre-treatment biopsy samples were profiled using the CosMx SMI platform. Initial data processing was conducted in Seurat v5.0^55^. Cells with fewer than 20 transcripts per cell or negative and system probe percentages greater than 10% were excluded from further analysis. FOVs were analyzed to identify biased gene expression profiles which were flagged for exclusion in downstream analyses and visualizations^56^. Samples were merged into a single Seurat object where they were log normalized and scaled followed by computing the principal component analysis reduction. Following initial dimensionality reduction, batch correction was conducted using the Harmony R package^57^. The batch corrected reduction was then used to generate UMAPs for cell visualization.

### Cell Annotation

Major cell type labels were assigned using the InSituTree R package^58^. Here, TNBC scRNA-seq reference cell profiles were curated from Wu et al^31^. Initially, 12 unique major cell types were identified where two cell types, Myoepithelial and T-lymphoid, were selected for more granular cell typing. CD4+ T cells and CD8+ T cells were distinguished using literature curated marker genes in the HieraType R package^59^. Cells labeled as myoepithelial were refined by recomputing the batch-corrected dimensionality reduction and sub clustering of cells. Differentially expressed genes among sub-clusters were identified using the ‘FindAllMarkers()’ function constrained to genes that were not prone to segmentation issues identified using the ‘overlap_ratio_metric()’ function in the smiDE R package^60^. Cell clusters with significant EMT-related gene expression were relabeled as mesenchymal-like cells while other clusters remained as myoepithelial. 15 final cell types were used for downstream analysis. Cell type proportion associations with treatment response were assessed using a permutation-based approach implemented in scProportionTest^61^.

### Niche Analysis

Global cellular niches were derived from local cellular neighborhood composition. Cellular neighborhoods were defined as cells found within a 15 µm radius from each cell’s centroid. Local neighborhoods composition profiles were compiled across all cells in all four samples. Recurring spatial niches were then identified using K-means clustering. The optimal number of neighborhoods identified were guided by inspection of the within-cluster sum of squares across additional K-clusters and further refined for interpretability. Seven final niches were identified reflecting unique spatial domains. Differential gene expression was assessed using DESeq2^62^. Raw gene expression of single cells was aggregated into pseudobulk profiles for each sample-niche combination. Differential expression was first modeled using a one-versus-rest approach with sample and niche mixed effects. Pairwise analysis of epithelial dominant niches 6 and 7 were also assessed using DESeq2. The Wald test statistics were used to rank genes for gene set enrichment analysis of niches using fgsea^9^. Enrichment was assessed using Hallmark gene sets retrieved from msigdb to summarize transcriptional programs in each niche^51^. Niche proportion associations with treatment response were assessed using scProportionTest^58^.

### Colocalization Network Analysis

Samples were modeled as marked spatial point patterns where cells were represented as points given segmented cell centroids and annotation labels. Bivariate cross pair correlation functions were applied between each cell type pair found in individual samples with translation edge correction using the SpatStat R package^63^. Statistical significance was determined from Monte Carlo envelopes by random relabeling cell types in 499 simulations. Z-scores were derived by assessing observed curves relative to the null distribution and summarized within a 0-50µm range. P-values were adjusted for multiple hypothesis testing using the Benjamini-Hochberg procedure. Statistically significant positive was visualized in network diagrams for individual samples using R package Igraph^64^. Colocalization scores across multiple samples were combined using Stouffer’s method. Consensus colocalization networks were constructed to represent global spatial relationships of enrichment. Colocalization networks were also constructed for individual samples.

Node-level summary metrics were computed within Igraph to characterize influence and colocalization trends between nodes. Overall colocalization strength was defined as the sum of edge weights connected to a given node. Average colocalization strength was derived by dividing the overall colocalization strength by the number of edges or degree of a given node. Interaction diversity was determined using Shannon entropy applied to normalized edge weights^65^. Here, higher values suggest even colocalization with multiple other nodes/cells while lower values indicate selective colocalization with few neighbor types. Eigenvector centrality characterizes the influence of nodes within the network. More specifically, higher eigenvector values indicate that nodes are connected to other highly influential or well-connected nodes. All scores were converted to z-scores for visualization.

Network communities were identified using the Infomap algorithm in Igraph^64–66^. Edges were reclassified as within- and between-community edges for calculation of summary metrics for each community. Community-community interactions were summarized by summing edge weights between community pairs. Community edge interactions were visualized in a heatmap where rows indicate the fraction of outgoing influence from each community to all communities. Community centralization was computed as the standard deviation of node-level betweenness centrality, reflecting the degree to which communities are dominated by central hub nodes. Community cohesion was as the sum of internal edge weights divided by the squared number of nodes in the community. Cohesion indicates the density and strength of connectivity of nodes within a community. Community polarity assesses asymmetry in spatial dependence of communities. It was defined as the fraction of mean outgoing edge weight relative to the total mean incoming and outgoing edge weights. Positive values indicate communities tend to colocalize with other communities while negative values suggest a community is spatially anchored and characterized relative to other communities.

## ACKNOWLEDGEMENTS

“Research reported in this publication was supported in part by the Emory University Integrated Genomics Core (EIGC; RRID:SCR_023529) of the Winship Cancer Institute of Emory University and NIH/NCI under award number, 2P30CA138292-04. The content is solely the responsibility of the authors and does not necessarily reflect the official views of the National Institute of Health.”

## REFERENCES

(1) Partridge, A. H.; Rumble, R. B.; Carey, L. A.; Come, S. E.; Davidson, N. E.; Di Leo, A.; Gralow, J.; Hortobagyi, G. N.; Moy, B.; Yee, D.;, et al. Chemotherapy and targeted therapy for women with human epidermal growth factor receptor 2-negative (or unknown) advanced breast cancer: American Society of Clinical Oncology Clinical Practice Guideline. J Clin Oncol 2014, 32 (29), 3307–3329. DOI: 10.1200/JCO.2014.56.7479.

(2) Mittendorf, E. A.; Zhang, H.; Barrios, C. H.; Saji, S.; Jung, K. H.; Hegg, R.; Koehler, A.; Sohn, J.; Iwata, H.; Telli, M. L.;, et al. Neoadjuvant atezolizumab in combination with sequential nab-paclitaxel and anthracycline-based chemotherapy versus placebo and chemotherapy in patients with early-stage triple-negative breast cancer (IMpassion031): a randomised, double-blind, phase 3 trial. Lancet 2020, 396 (10257), 1090–1100. DOI: 10.1016/S0140-6736(20)31953-X.

(3) Gianni, L.; Huang, C. S.; Egle, D.; Bermejo, B.; Zamagni, C.; Thill, M.; Anton, A.; Zambelli, S.; Bianchini, G.; Russo, S.;, et al. Pathologic complete response (pCR) to neoadjuvant treatment with or without atezolizumab in triple-negative, early high-risk and locally advanced breast cancer: NeoTRIP Michelangelo randomized study. Ann Oncol 2022, 33 (5), 534–543. DOI: 10.1016/j.annonc.2022.02.004.

(4) Schmid, P.; Cortes, J.; Pusztai, L.; McArthur, H.; Kummel, S.; Bergh, J.; Denkert, C.; Park, Y. H.; Hui, R.; Harbeck, N.;, et al. Pembrolizumab for Early Triple-Negative Breast Cancer. N Engl J Med 2020, 382 (9), 810–821. DOI: 10.1056/NEJMoa1910549.

(5) Goncalves, A.; de Nonneville, A. Tailoring chemoimmunotherapy de-escalation in early-stage triple-negative breast cancer. Lancet Oncol 2025, 26 (3), 273–274. DOI: 10.1016/S1470-2045(25)00028-2.

(6) Carvalho, F. M. Targeting low-risk triple-negative breast cancer: a review on de-escalation strategies for a new era. Transl Breast Cancer Res 2025, 6, 4. DOI: 10.21037/tbcr-24-28.

(7) Gupta, R. K.; Roy, A. M.; Gupta, A.; Takabe, K.; Dhakal, A.; Opyrchal, M.; Kalinski, P.; Gandhi, S. Systemic Therapy De-Escalation in Early-Stage Triple-Negative Breast Cancer: Dawn of a New Era? Cancers (Basel) 2022, 14 (8). DOI: 10.3390/cancers14081856.

(8) Wood, S. J.; Gao, Y.; Lee, J.-H.; Chen, J.; Wang, Q.; Meisel, J. L.; Li, X. High tumor infiltrating lymphocytes are significantly associated with pathological complete response in triple negative breast cancer treated with neoadjuvant KEYNOTE-522 chemoimmunotherapy. Breast Cancer Research and Treatment 2024, 205 (1), 193–199. DOI: 10.1007/s10549-023-07233-2.

(9) Bianchini, G.; De Angelis, C.; Licata, L.; Gianni, L. Treatment landscape of triple-negative breast cancer - expanded options, evolving needs. Nat Rev Clin Oncol 2022, 19 (2), 91–113. DOI: 10.1038/s41571-021-00565-2.

(10) Lehmann, B. D.; Bauer, J. A.; Chen, X.; Sanders, M. E.; Chakravarthy, A. B.; Shyr, Y.; Pietenpol, J. A. Identification of human triple-negative breast cancer subtypes and preclinical models for selection of targeted therapies. J Clin Invest 2011, 121 (7), 2750–2767. DOI: 10.1172/JCI45014.

(11) Yin, L.; Duan, J. J.; Bian, X. W.; Yu, S. C. Triple-negative breast cancer molecular subtyping and treatment progress. Breast Cancer Res 2020, 22 (1), 61. DOI: 10.1186/s13058-020-01296-5.

(12) Wolf, D. M.; Yau, C.; Wulfkuhle, J.; Brown-Swigart, L.; Gallagher, R. I.; Lee, P. R. E.; Zhu, Z.; Magbanua, M. J.; Sayaman, R.; O’Grady, N.;, et al. Redefining breast cancer subtypes to guide treatment prioritization and maximize response: Predictive biomarkers across 10 cancer therapies. Cancer Cell 2022, 40 (6), 609–623 e606. DOI: 10.1016/j.ccell.2022.05.005.

(13) Lehmann, B. D.; Jovanovic, B.; Chen, X.; Estrada, M. V.; Johnson, K. N.; Shyr, Y.; Moses, H. L.; Sanders, M. E.; Pietenpol, J. A. Refinement of Triple-Negative Breast Cancer Molecular Subtypes: Implications for Neoadjuvant Chemotherapy Selection. PLoS One 2016, 11 (6), e0157368. DOI: 10.1371/journal.pone.0157368.

(14) Paik, S.; Shak, S.; Tang, G.; Kim, C.; Baker, J.; Cronin, M.; Baehner, F. L.; Walker, M. G.; Watson, D.; Park, T.;, et al. A multigene assay to predict recurrence of tamoxifen-treated, node-negative breast cancer. N Engl J Med 2004, 351 (27), 2817–2826. DOI: 10.1056/NEJMoa041588.

(15) Ross, J. S.; Hatzis, C.; Symmans, W. F.; Pusztai, L.; Hortobagyi, G. N. Commercialized multigene predictors of clinical outcome for breast cancer. Oncologist 2008, 13 (5), 477–493. DOI: 10.1634/theoncologist.2007-0248.

(16) Pennock, N. D.; Jindal, S.; Horton, W.; Sun, D.; Narasimhan, J.; Carbone, L.; Fei, S. S.; Searles, R.; Harrington, C. A.; Burchard, J.;, et al. RNA-seq from archival FFPE breast cancer samples: molecular pathway fidelity and novel discovery. BMC Med Genomics 2019, 12 (1), 195. DOI: 10.1186/s12920-019-0643-z.

(17) Andre, F.; Ismaila, N.; Henry, N. L.; Somerfield, M. R.; Bast, R. C.; Barlow, W.; Collyar, D. E.; Hammond, M. E.; Kuderer, N. M.; Liu, M. C.;, et al. Use of Biomarkers to Guide Decisions on Adjuvant Systemic Therapy for Women With Early-Stage Invasive Breast Cancer: ASCO Clinical Practice Guideline Update-Integration of Results From TAILORx. J Clin Oncol 2019, 37 (22), 1956–1964. DOI: 10.1200/JCO.19.00945.

(18) Stover, D. G.; Coloff, J. L.; Barry, W. T.; Brugge, J. S.; Winer, E. P.; Selfors, L. M. The Role of Proliferation in Determining Response to Neoadjuvant Chemotherapy in Breast Cancer: A Gene Expression-Based Meta-Analysis. Clin Cancer Res 2016, 22 (24), 6039–6050. DOI: 10.1158/1078-0432.CCR-16-0471.

(19) Chen, X.; Li, J.; Gray, W. H.; Lehmann, B. D.; Bauer, J. A.; Shyr, Y.; Pietenpol, J. A. TNBCtype: A Subtyping Tool for Triple-Negative Breast Cancer. Cancer Inform 2012, 11, 147–156. DOI: 10.4137/CIN.S9983.

(20) Su, S.; Chen, J.; Yao, H.; Liu, J.; Yu, S.; Lao, L.; Wang, M.; Luo, M.; Xing, Y.; Chen, F.;, et al. CD10(+)GPR77(+) Cancer-Associated Fibroblasts Promote Cancer Formation and Chemoresistance by Sustaining Cancer Stemness. Cell 2018, 172 (4), 841–856 e816. DOI: 10.1016/j.cell.2018.01.009.

(21) Wu, S. Z.; Roden, D. L.; Wang, C.; Holliday, H.; Harvey, K.; Cazet, A. S.; Murphy, K. J.; Pereira, B.; Al-Eryani, G.; Bartonicek, N.;, et al. Stromal cell diversity associated with immune evasion in human triple-negative breast cancer. EMBO J 2020, 39 (19), e104063. DOI: 10.15252/embj.2019104063.

(22) Arole, V.; Nitta, H.; Wei, L.; Shen, T.; Parwani, A. V.; Li, Z. M2 tumor-associated macrophages play important role in predicting response to neoadjuvant chemotherapy in triple-negative breast carcinoma. Breast Cancer Res Treat 2021, 188 (1), 37–42. DOI: 10.1007/s10549-021-06260-1.

(23) Alizadeh, D.; Trad, M.; Hanke, N. T.; Larmonier, C. B.; Janikashvili, N.; Bonnotte, B.; Katsanis, E.; Larmonier, N. Doxorubicin eliminates myeloid-derived suppressor cells and enhances the efficacy of adoptive T-cell transfer in breast cancer. Cancer Res 2014, 74 (1), 104–118. DOI: 10.1158/0008-5472.CAN-13-1545.

(24) Kashima, H.; Momose, F.; Umehara, H.; Miyoshi, N.; Ogo, N.; Muraoka, D.; Shiku, H.; Harada, N.; Asai, A. Epirubicin, Identified Using a Novel Luciferase Reporter Assay for Foxp3 Inhibitors, Inhibits Regulatory T Cell Activity. PLoS One 2016, 11 (6), e0156643. DOI: 10.1371/journal.pone.0156643.

(25) Helwick, C. Demystifying Immunotherapy for Early-Stage Triple-Negative Breast Cancer. The ASCO Post, 2022.

(26) Chen, B.; Khodadoust, M. S.; Liu, C. L.; Newman, A. M.; Alizadeh, A. A. Profiling Tumor Infiltrating Immune Cells with CIBERSORT. Methods Mol Biol 2018, 1711, 243–259. DOI: 10.1007/978-1-4939-7493-1_12.

(27) Park, S.; Ock, C. Y.; Kim, H.; Pereira, S.; Park, S.; Ma, M.; Choi, S.; Kim, S.; Shin, S.; Aum, B. J.;, et al. Artificial Intelligence-Powered Spatial Analysis of Tumor-Infiltrating Lymphocytes as Complementary Biomarker for Immune Checkpoint Inhibition in Non-Small-Cell Lung Cancer. J Clin Oncol 2022, 40 (17), 1916–1928. DOI: 10.1200/JCO.21.02010.

(28) Ali, H. R.; Chlon, L.; Pharoah, P. D.; Markowetz, F.; Caldas, C. Patterns of Immune Infiltration in Breast Cancer and Their Clinical Implications: A Gene-Expression-Based Retrospective Study. PLoS Med 2016, 13 (12), e1002194. DOI: 10.1371/journal.pmed.1002194.

(29) Craven, K. E.; Gokmen-Polar, Y.; Badve, S. S. CIBERSORT analysis of TCGA and METABRIC identifies subgroups with better outcomes in triple negative breast cancer. Sci Rep 2021, 11 (1), 4691. DOI: 10.1038/s41598-021-83913-7.

(30) Kim, C.; Gao, R.; Sei, E.; Brandt, R.; Hartman, J.; Hatschek, T.; Crosetto, N.; Foukakis, T.; Navin, N. E. Chemoresistance Evolution in Triple-Negative Breast Cancer Delineated by Single-Cell Sequencing. Cell 2018, 173 (4), 879–893 e813. DOI: 10.1016/j.cell.2018.03.041.

(31) Wu, S. Z.; Al-Eryani, G.; Roden, D. L.; Junankar, S.; Harvey, K.; Andersson, A.; Thennavan, A.; Wang, C.; Torpy, J. R.; Bartonicek, N.;, et al. A single-cell and spatially resolved atlas of human breast cancers. Nat Genet 2021, 53 (9), 1334–1347. DOI: 10.1038/s41588-021-00911-1.

(32) Jayasingam, S. D.; Citartan, M.; Thang, T. H.; Zin, A. A. M.; Ang, K. C.; Ch’ng, E. S. Evaluating the Polarization of Tumor-Associated Macrophages Into M1 and M2 Phenotypes in Human Cancer Tissue: Technicalities and Challenges in Routine Clinical Practice. Front Oncol 2020, 9. DOI: ARTN 1512 10.3389/fonc.2019.01512.

(33) Tiainen, S.; Tumelius, R.; Rilla, K.; Hämäläinen, K.; Tammi, M.; Tammi, R.; Kosma, V. M.; Oikari, S.; Auvinen, P. High numbers of macrophages, especially M2-like (CD163-positive), correlate with hyaluronan accumulation and poor outcome in breast cancer. Histopathology 2015, 66 (6), 873– 883. DOI: 10.1111/his.12607.

(34) RStudio: Integrated Development for R; RStudio: 2020.

(35) Cancer Genome Atlas Research, N.; Weinstein, J. N.; Collisson, E. A.; Mills, G. B.; Shaw, K. R.; Ozenberger, B. A.; Ellrott, K.; Shmulevich, I.; Sander, C.; Stuart, J. M. The Cancer Genome Atlas Pan-Cancer analysis project. Nat Genet 2013, 45 (10), 1113–1120. DOI: 10.1038/ng.2764.

(36) Hoadley, K. A.; Yau, C.; Hinoue, T.; Wolf, D. M.; Lazar, A. J.; Drill, E.; Shen, R.; Taylor, A. M.; Cherniack, A. D.; Thorsson, V.;, et al. Cell-of-Origin Patterns Dominate the Molecular Classification of 10,000 Tumors from 33 Types of Cancer. Cell 2018, 173 (2), 291–304 e296. DOI: 10.1016/j.cell.2018.03.022.

(37) Hatzis, C.; Pusztai, L.; Valero, V.; Booser, D. J.; Esserman, L.; Lluch, A.; Vidaurre, T.; Holmes, F.; Souchon, E.; Wang, H.;, et al. A genomic predictor of response and survival following taxane-anthracycline chemotherapy for invasive breast cancer. JAMA 2011, 305 (18), 1873–1881. DOI: 10.1001/jama.2011.593.

(38) Horak, C. E.; Pusztai, L.; Xing, G.; Trifan, O. C.; Saura, C.; Tseng, L. M.; Chan, S.; Welcher, R.; Liu, D. Biomarker analysis of neoadjuvant doxorubicin/cyclophosphamide followed by ixabepilone or Paclitaxel in early-stage breast cancer. Clin Cancer Res 2013, 19 (6), 1587–1595. DOI: 10.1158/1078-0432.CCR-12-1359.

(39) Gluck, S.; Ross, J. S.; Royce, M.; McKenna, E. F., Jr.; Perou, C. M.; Avisar, E.; Wu, L. TP53 genomics predict higher clinical and pathologic tumor response in operable early-stage breast cancer treated with docetaxel-capecitabine +/-trastuzumab. Breast Cancer Res Treat 2012, 132 (3), 781–791. DOI: 10.1007/s10549-011-1412-7.

(40) Esserman, L. J.; Berry, D. A.; Cheang, M. C.; Yau, C.; Perou, C. M.; Carey, L.; DeMichele, A.; Gray, J. W.; Conway-Dorsey, K.; Lenburg, M. E.;, et al. Chemotherapy response and recurrence-free survival in neoadjuvant breast cancer depends on biomarker profiles: results from the I-SPY 1 TRIAL (CALGB 150007/150012; ACRIN 6657). Breast Cancer Res Treat 2012, 132 (3), 1049–1062. DOI: 10.1007/s10549-011-1895-2.

(41) Miyake, T.; Nakayama, T.; Naoi, Y.; Yamamoto, N.; Otani, Y.; Kim, S. J.; Shimazu, K.; Shimomura, A.; Maruyama, N.; Tamaki, Y.;, et al. GSTP1 expression predicts poor pathological complete response to neoadjuvant chemotherapy in ER-negative breast cancer. Cancer Sci 2012, 103 (5), 913–920. DOI: 10.1111/j.1349-7006.2012.02231.x.

(42) Bray, N. L.; Pimentel, H.; Melsted, P.; Pachter, L. Near-optimal probabilistic RNA-seq quantification. Nat Biotechnol 2016, 34 (5), 525–527. DOI: 10.1038/nbt.3519.

(43) Karaayvaz, M.; Cristea, S.; Gillespie, S. M.; Patel, A. P.; Mylvaganam, R.; Luo, C. C.; Specht, M. C.; Bernstein, B. E.; Michor, F.; Ellisen, L. W. Unravelling subclonal heterogeneity and aggressive disease states in TNBC through single-cell RNA-seq. Nat Commun 2018, 9 (1), 3588. DOI: 10.1038/s41467-018-06052-0.

(44) Alvarez, M. J.; Shen, Y.; Giorgi, F. M.; Lachmann, A.; Ding, B. B.; Ye, B. H.; Califano, A. Functional characterization of somatic mutations in cancer using network-based inference of protein activity. Nat Genet 2016, 48 (8), 838–847. DOI: 10.1038/ng.3593.

(45) rms: Regression Modeling Strategies; The Comprehensive R Archive Network: https://CRAN.R-project.org/package=rms, 2023.

(46) survminer: Drawing Survival Curves using ’ggplot2’; The Comprehensive R Archive Network: https://CRAN.R-project.org/package=survminer, 2021.

(47) gplots: Various R Programming Tools for Plotting Data; The Comprehensive R Archive Network: https://CRAN.R-project.org/package=gplots, 2022.

(48) RColorBrewer: ColorBrewer Palettes; https://CRAN.R-project.org/package=RColorBrewer: 2022.

(49) R: A Language and Environment for Statistical Computing; R Foundation for Statistical Computing: https://www.R-project.org/, 2023.

(50) Larsson J; P, G. A Case Study in Fitting Area-Proportional Euler Diagrams with Ellipses Using eulerr . Proceedings of International Workshop on Set Visualization and Reasoning 2018, 2116, 84–91.

(51) msigdbr: MSigDB Gene Sets for Multiple Organisms in a Tidy Data Format; The Comprehensive R Archive Network: https://CRAN.R-project.org/package=msigdbr, 2022.

(52) Liberzon, A.; Birger, C.; Thorvaldsdottir, H.; Ghandi, M.; Mesirov, J. P.; Tamayo, P. The Molecular Signatures Database (MSigDB) hallmark gene set collection. Cell Syst 2015, 1 (6), 417–425. DOI: 10.1016/j.cels.2015.12.004.

(53) Moffitt, J. R.; Hao, J.; Wang, G.; Chen, K. H.; Babcock, H. P.; Zhuang, X. High-throughput single-cell gene-expression profiling with multiplexed error-robust fluorescence in situ hybridization. Proceedings of the National Academy of Sciences 2016, 113 (39), 11046–11051. DOI: 10.1073/pnas.1612826113 (accessed 2026/05/07).

(54) He, S.; Bhatt, R.; Brown, C.; Brown, E. A.; Buhr, D. L.; Chantranuvatana, K.; Danaher, P.; Dunaway, D.; Garrison, R. G.; Geiss, G.;, et al. High-plex imaging of RNA and proteins at subcellular resolution in fixed tissue by spatial molecular imaging. Nat Biotechnol 2022, 40 (12), 1794–1806. DOI: 10.1038/s41587-022-01483-z.

(55) Hao, Y.; Stuart, T.; Kowalski, M. H.; Choudhary, S.; Hoffman, P.; Hartman, A.; Srivastava, A.; Molla, G.; Madad, S.; Fernandez-Granda, C.;, et al. Dictionary learning for integrative, multimodal and scalable single-cell analysis. Nat Biotechnol 2024, 42 (2), 293–304. DOI: 10.1038/s41587-023-01767-y From NLM Medline.

(56) CosMx-Analysis-Scratch-Space; GitHub: GitHub, 2026. https://github.com/Nanostring-Biostats/CosMx-Analysis-Scratch-Space (accessed 2/23/26).

(57) Korsunsky, I.; Millard, N.; Fan, J.; Slowikowski, K.; Zhang, F.; Wei, K.; Baglaenko, Y.; Brenner, M.; Loh, P. R.; Raychaudhuri, S. Fast, sensitive and accurate integration of single-cell data with Harmony. Nat Methods 2019, 16 (12), 1289–1296. DOI: 10.1038/s41592-019-0619-0 From NLM Medline.

(58) InSituTree: Run Hierarchical Cell Typing; GitHub: GitHub, 2025. https://github.com/Nanostring-Biostats/CosMx-Analysis-Scratch-Space/_code/InSituTree.

(59) HieraType: Package for Hierarchical Cell Typing using Smoothed Metagene Scores; GitHub: GitHub, 2025. https://github.com/Nanostring-Biostats/CosMx-Analysis-Scratch-Space/_code/HieraType.

(60) Vasconcelos, A. G.; McGuire, D.; Simon, N.; Danaher, P.; Shojaie, A. Differential Expression Analysis for Spatially Correlated Data. BioRxiv 2025.DOI: 10.1101/2024.08.02.606405.

(61) Miller, S. A.; Policastro, R. A.; Sriramkumar, S.; Lai, T.; Huntington, T. D.; Ladaika, C. A.; Kim, D.; Hao, C.; Zentner, G. E.; O’Hagan, H. M. LSD1 and Aberrant DNA Methylation Mediate Persistence of Enteroendocrine Progenitors That Support BRAF-Mutant Colorectal Cancer. Cancer Res 2021, 81 (14), 3791–3805. DOI: 10.1158/0008-5472.CAN-20-3562 From NLM Medline.

(62) Love, M. I.; Huber, W.; Anders, S. Moderated estimation of fold change and dispersion for RNA-seq data with DESeq2. Genome Biol 2014, 15 (12), 550. DOI: 10.1186/s13059-014-0550-8 From NLM Medline.

(63) Baddeley, A.; Rubuk, E.; Turner, R. Spatial Point Patterns: Methodology and Applications with R; Chapman and Hall/CRC Press,London. , 2015.

(64) Csárdi G, N. T. igraph: Network Analysis and Visualization in R. *InterJournal*, Complex Systems 2006, 1685.

(65) Freitas, C. G. S.; Aquino, A. L. L.; Ramos, H. S.; Frery, A. C.; Rosso, O. A. A detailed characterization of complex networks using Information Theory. Sci Rep 2019, 9 (1), 16689. DOI: 10.1038/s41598-019-53167-5 From NLM PubMed-not-MEDLINE.

(66) Rosvall, M.; Bergstrom, C. T. Maps of random walks on complex networks reveal community structure. Preceeding of the National Academy of Sciences 2008, 105 (4).

