## Supplemental Figures for "Spatial Logic Reconciles Gene-signature Methods in Triple Negative Breast Cancer"

SUPPLEMENTAL MATERIAL

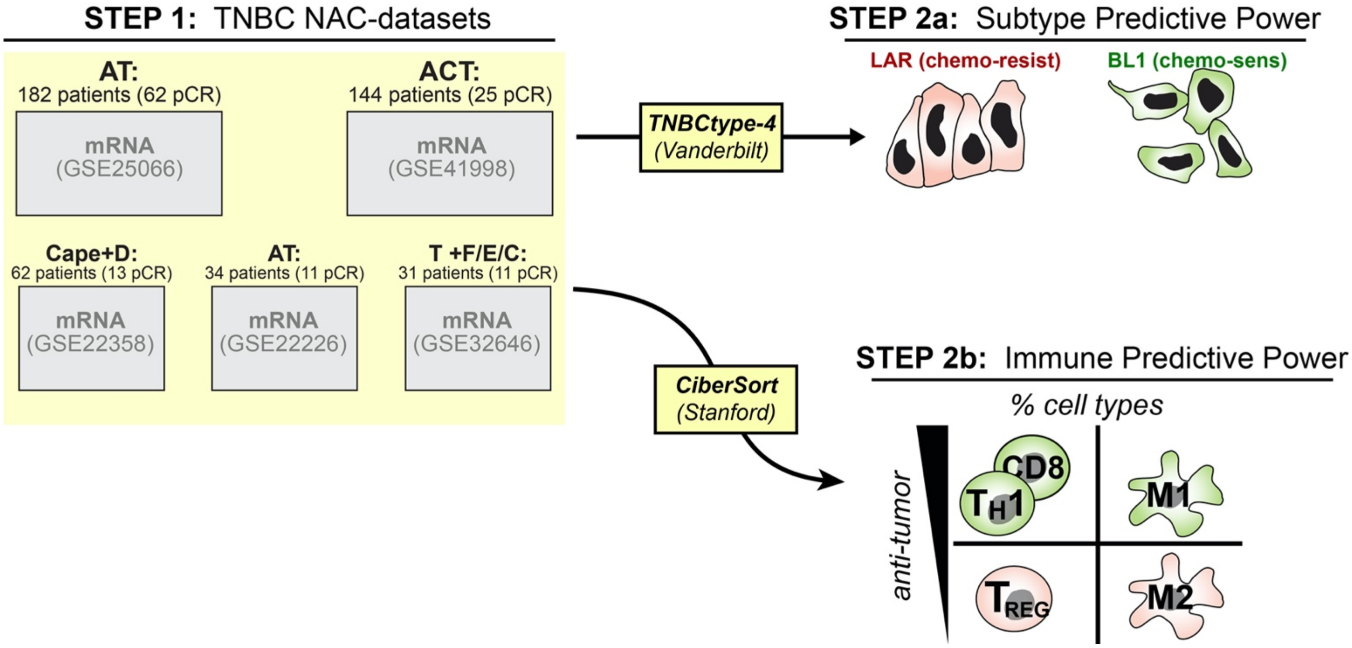


**Figure S01. Breast cancer-subtype and infiltrating macrophage signatures predict chemotherapy-response and survival.** Graphic Overview of NAC datasets used for TNBC-type4 development and validation. These datasets were used in this study to benchmark TNBC-type and CIBERSORT predictive power (Normalized signature Scores are available in Supporting Table #

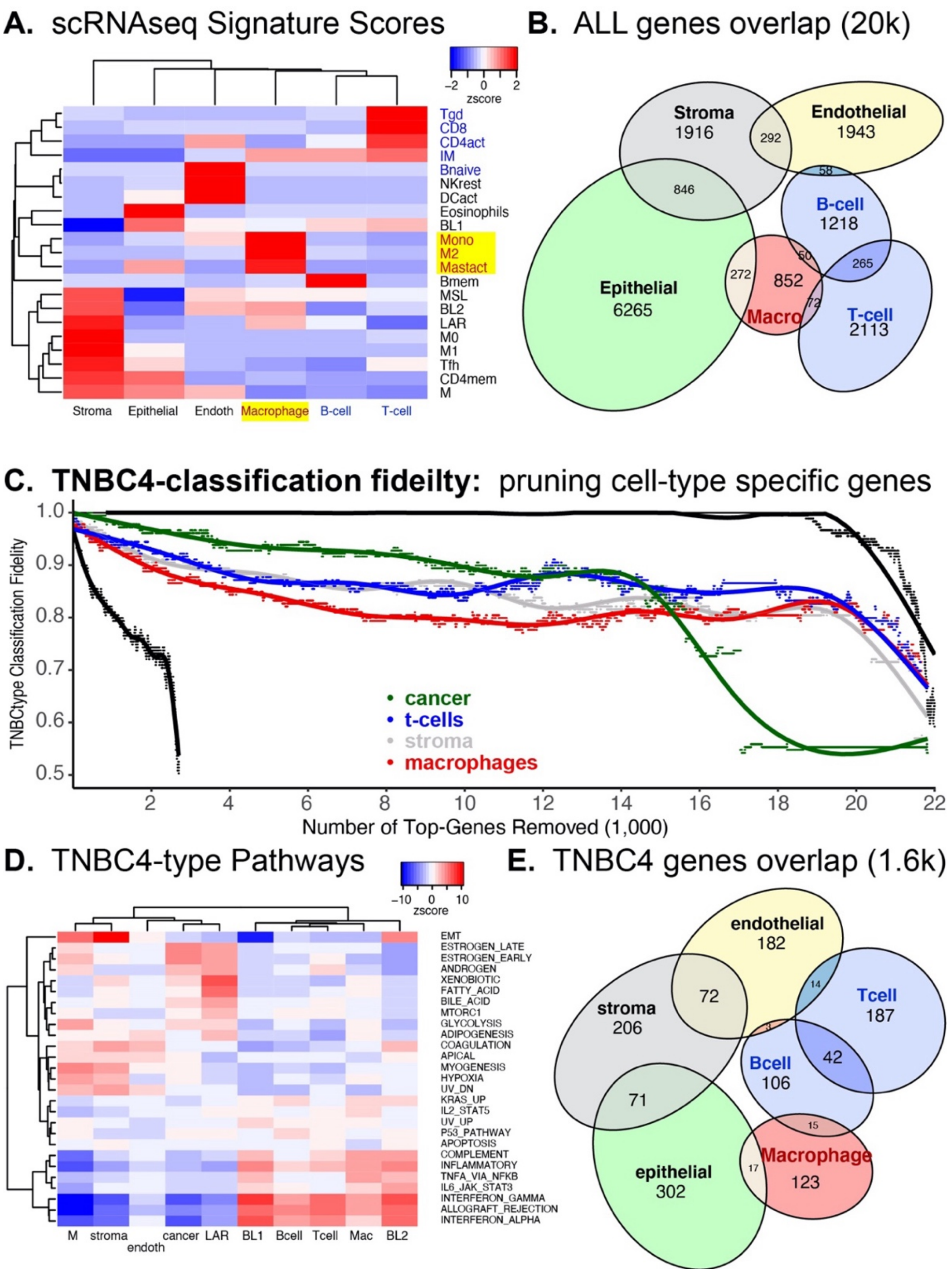


**Figure S02. Comparison of CIBERSORT and TNBC-type signature enrichment across single-cell RNAseq Datasets A.** Enrichment scores (z-score) of CIBERSORT and TNBC-type on scRNAseq dataset published for breast cancer in 2018. **B.** Cell-type specific expression across breast tumor cell-compartments demonstrates independence and overlap of genes. **C.** Evaluation of TNBC-classification fidelity with removal of top features/genes from non-cancer-cell-types that often infiltrate tumors. **D.** Evaluation of CIBERSORT-t-cell fidelity with removal of top features/genes from different cell types within tumors.

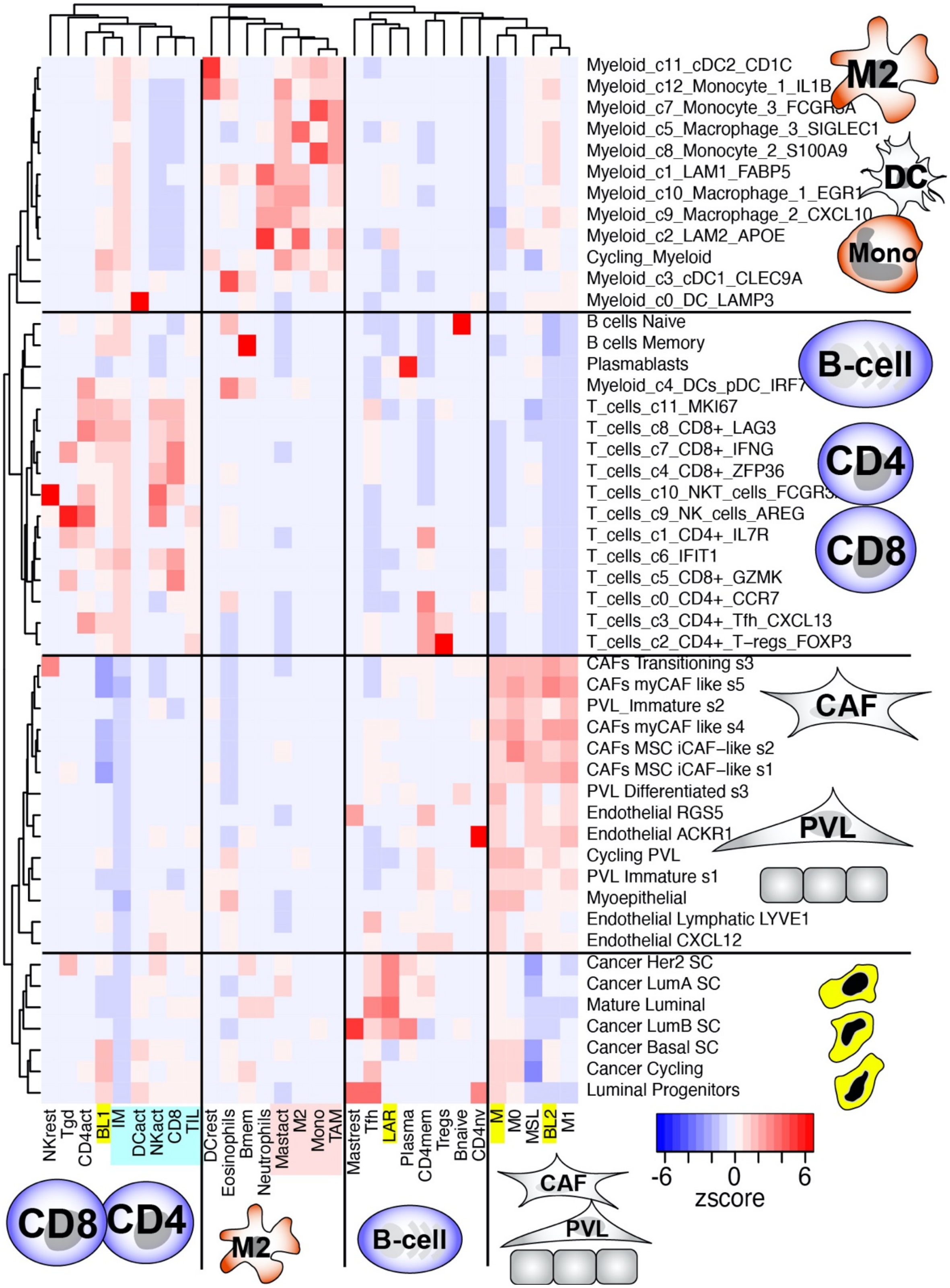


**Figure S03.** CIBERSORT and TNBC-type signature enrichment across 2021 breast tumor scRNAseq Atlas (GSE176078)

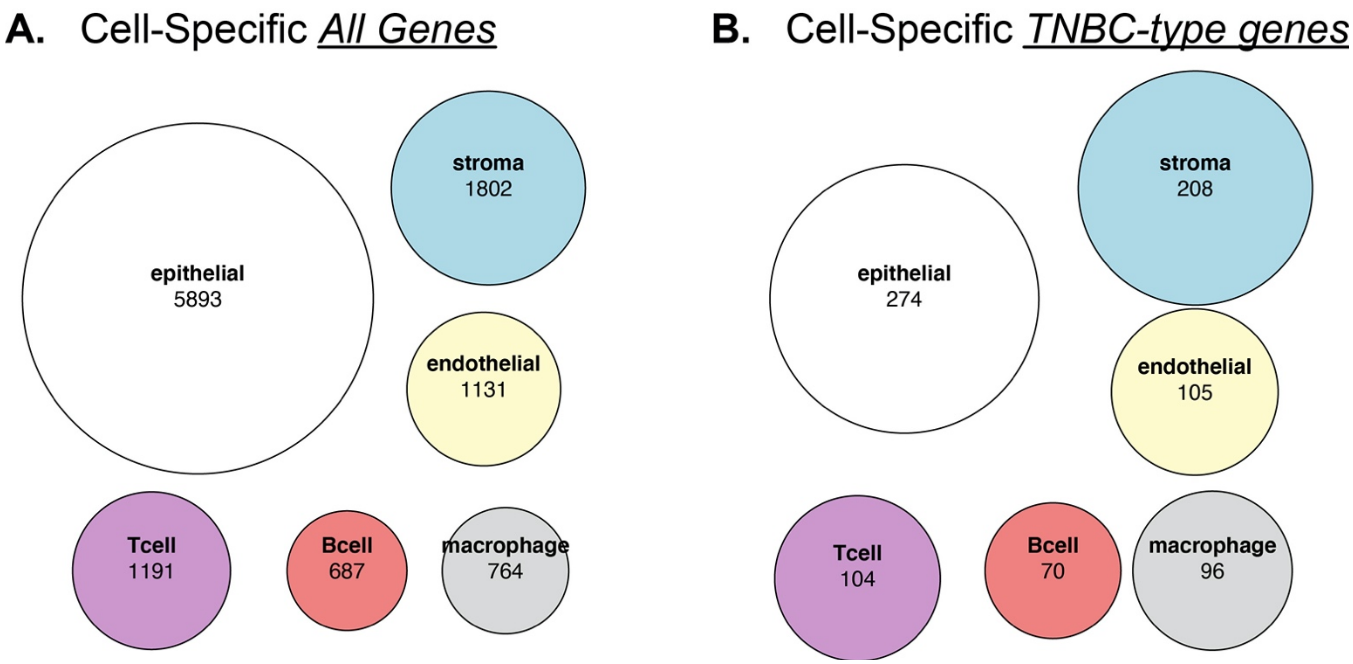


**Figure S04.** A. Cell-type specific expression of all protein-coding genes within scRNAseq data set (GSE118389) B. Cell-type selective expression of the top 1600 TNBC-type genes (GSE118389).

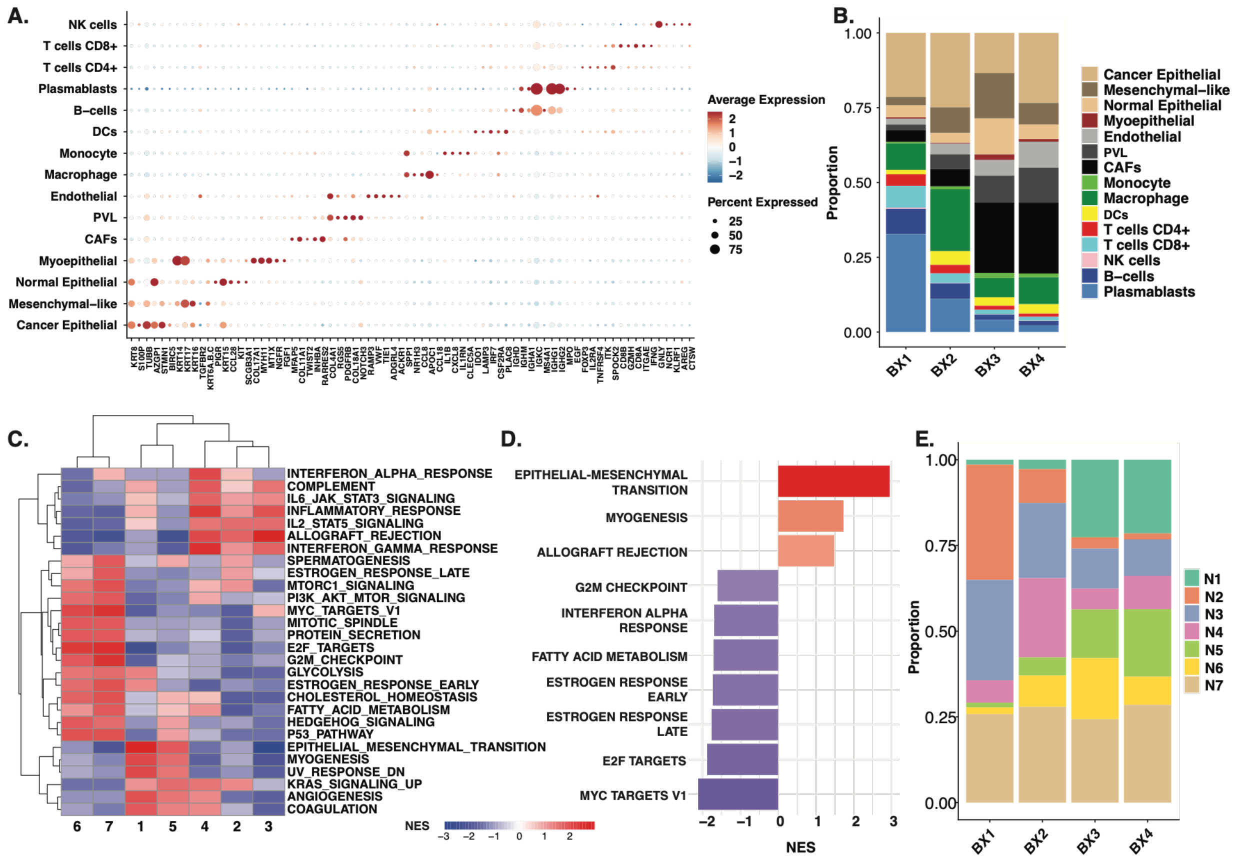


**Figure S05. Cell annotation and niche identification validation. A)** Top 5 differentially expressed markers for each cell population across all 4 spatial samples. **B)** Cell type proportions per sample. Samples BX1 and BX2 represent pCR samples, while BX3 and BX4 were no pCR samples. **C.** Hallmark pathway enrichment of the 7 niches with differentially expressed genes derived from a one-vs-rest model design using DESeq2. **D.** Hallmark pathways enriched in niche 6 relative to niche 7 demonstrating the upregulation of epithelial-mesenchymal transition and myogenesis pathways. **E.** Niche proportions per sample.

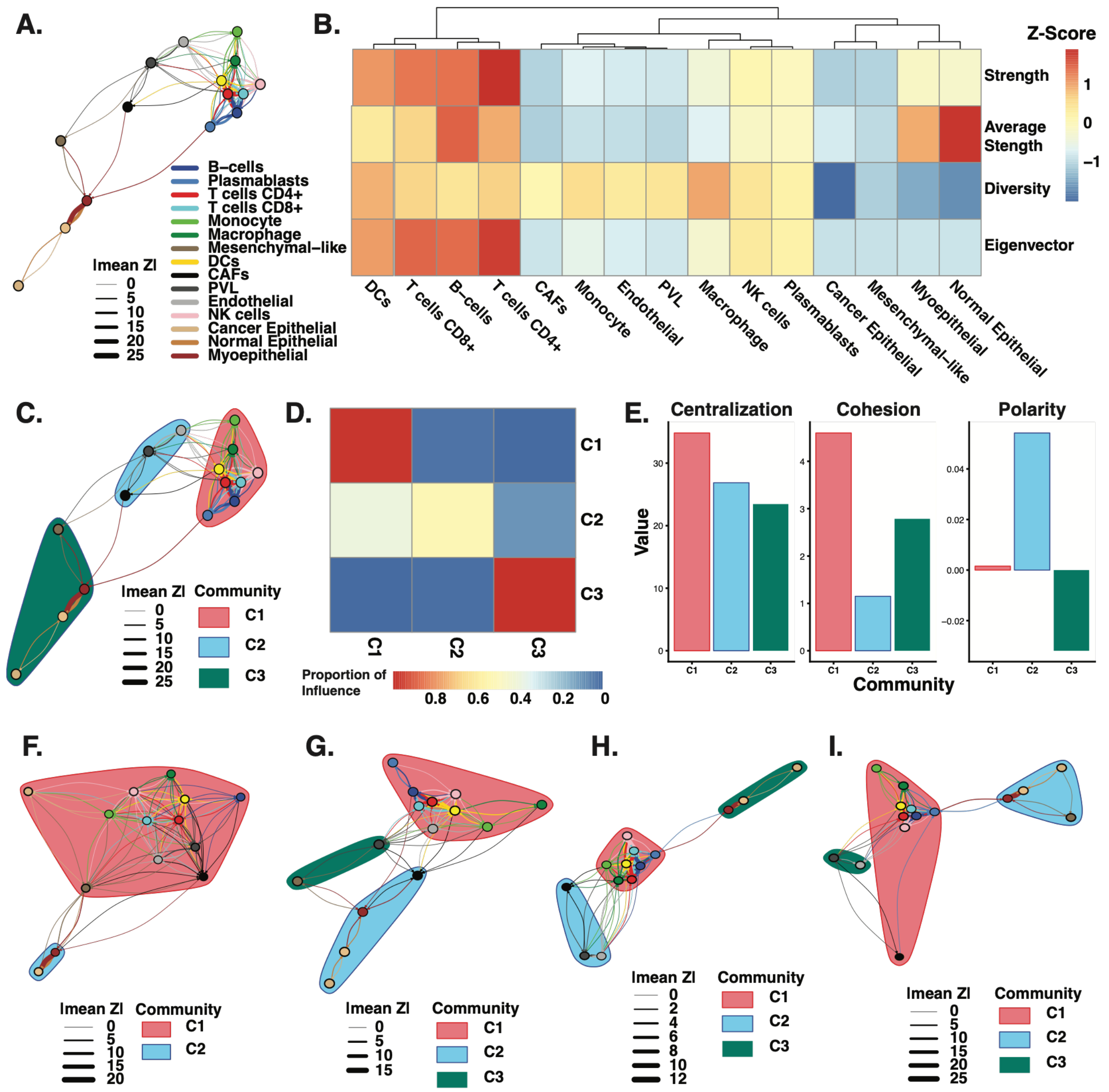


**Figure S06 Colocalization network analysis. A)** Consensus colocalization network derived from cross-pcf scores across all cell type pairs. Edges represent statistically significant colocalization between cell types where weight corresponds to the mean Z-score across samples. **B)** Node-level network metric z-scores including overall strength of interactions, average strength of interactions, diversity, and influence (eigenvector centrality). Notably, lymphocytes and DCs lead in strength and influence relative to other cell types. **C)** Consensus colocalization network annotated with communities detected using the Infomap community detection algorithm. **D)** Community-community colocalization heatmap. Values depict proportion of influence where rows sum to 1 and represent the proportion of interactions that form between other communities. **E)** Community validation through centralization, cohesion, and polarity metrics. High centralization indicated the presence of central or hub nodes within communities. Cohesion assesses the overall density and strength of interactions within communities. Polarity addresses the asymmetry of colocalization between communities. Positive polarity indicates communities that colocalize with nodes found in other communities while negative polarity indicates spatial organization is relative to other communities. **F-I)** Colocalization networks and communities of nodes in sample BX1, BX2, BX3, and BX4, respectively.
